# There is an “I” in team: individual improvements in supercharged cellulase cocktail facilitates cooperative cellulose degradation

**DOI:** 10.1101/2025.10.16.682856

**Authors:** Antonio DeChellis, Samantha Shimabukuro, Sumay Trivedi, Reena Lubowski, Bhargava Nemmaru, Shishir P. S. Chundawat

## Abstract

Lignocellulosic biomass is a vastly abundant renewable carbon source for biofuel production but its conversion to fermentable sugars is significantly hindered by an inherent recalcitrance to enzymatic degradation. Pretreatment technologies are successful in alleviating some challenges related to substrate recalcitrance, yet enzymes like cellulases still exhibit poor activity on highly crystalline and insoluble cellulose. Both cellulose and lignin present several issues with productive enzyme binding and efficient catalytic turnover. To address these bottlenecks, we employed protein supercharging to rationally design a glycosyl hydrolase (GH) family-6 exocellulase (Cel6B) and its native fused family-2a carbohydrate binding module (CBM2a) from the thermophilic cellulolytic microbe *Thermobifida fusca*. A total of 16 supercharged variants were designed across both GH/CBM domains and a chimeric library of 32 constructs, including the native enzyme, were synthesized and expressed in *E. coli*. The entire library of supercharged enzymes was tested for activity on several cellulosic substrates to identify one key construct, D5 CBM2a – WT Cel6B, that had a positively supercharged CBM2a that showed 2-3-fold higher activity on all substrates tested at pH 5.5. Purified enzyme assays confirmed that exocellulases behave quite different from their endocellulase counterparts when supercharged using similar protocols. Still, the purified D5 CBM2a – WT Cel6B mutant showed a 2.3-fold improvement in specific activity compared to native enzyme on crystalline cellulose. Analysis of melt curves depict that, while all other constructs tested have one distinct melt peak near the expected CBM melting point, domain melting is decoupled for the D5 CBM2a mutant. This effect reveals an intrinsic melting temperature of the Cel6B CD nearly 18 °C higher than the coupled melting temperature of the full-length enzyme. This unexpected catalytic domain stabilization effect of supercharged CBM2a domain is likely the driving force for activity improvements seen for this exocellulase that is otherwise prone to stalling and denaturation on the cellulose surface during processive catalytic turnover cycles. When combining this supercharged exocellulase construct with its endocellulase counterpart, our results show that supercharged enzymes that show the highest activity alone, produced the best synergistic partners. This study highlights another successful implementation of protein supercharging strategy for cellulases and provides another key piece towards building an effective synergistic cellulase cocktail for lignocellulosic biomass deconstruction.

## INTRODUCTION

Lignocellulosic biomass is the most widely available renewable energy source yet is underutilized due to its recalcitrance to microbial degradation.^1,2^ This carbon rich feedstock is readily sourced from agricultural and forestry wastes^3^ and is a rich source of polysaccharides like cellulose and hemicellulose which can be enzymatically saccharified by Carbohydrate Active enZymes (CAZymes) into soluble sugars and fermented into fuels like ethanol or butanol.^4,5^ The inherent recalcitrance towards enzymatic degradation is caused by several factors including; (i) limited access to cellulose and hemicellulose substrates within the core of plant cell walls, (ii) non-productive enzyme binding to structural phenolic polymer lignin and/or hemi/cellulose itself, and (iii) low activity on an insoluble and tightly packed crystalline structure of cellulose.^6–9^ This recalcitrance is partially overcome by thermochemical pretreatments that disrupt the overall biomass ultrastructure, redistribute lignin, and potentially alter cellulose crystallinity.^10^ Several technologies exist including alkaline methods that preserve hemicellulose like ammonium fiber expansion (AFEX) pretreatment that swells biomass fibers redistributing lignin and makes both hemicellulose and amorphous cellulose more readily accessible to enzymes.^11,12^ Extractive ammonia (EA) pretreatment builds on the original AFEX method to partially solubilize lignin and alter the crystalline structure of cellulose.^13^ One of the simplest, scalable, and most industrially mature technology is dilute acid (DA) pretreatment that relies on acids like sulfuric acid to hydrolyze and remove solubilized hemicellulose oligo-/mono-mers resulting in more accessible crystalline cellulose enriched with condensed lignin deposits still present.^14,15^ This is not a perfect solution as remaining lignin can still non-productively adsorb enzymes inhibiting its availability and/or activity, and pretreatment also doesn’t address challenges related to poor interfacial activity between a soluble enzyme and its insoluble crystalline substrate like cellulose.^16^ These factors significantly drive up overall processing costs for valorizing lignocellulosic wastes into valuable fuels and other bioproducts.^17,18^ To overcome these challenges, the inclusion of sacrificial proteins to block lignin binding and high enzyme loadings required to overcome low activity that significantly hinder the economic viability of overall process,^19–21^ thus rational design of better enzymes is needed.

To effectively hydrolyze the cellulose portion of biomass, a minimal cocktail comprising of at least an endocellulase, exocellulase, and glucosidase enzymes is required. These enzymes work together to synergistically cleave glycosidic linkages within the cellulose backbone down to cellobiose or cellobiosyl oligomers which is ultimately processed to glucose by a β-glucosidase.^4,22^ There are several industrially relevant enzymes available from different microbial hosts, and one such, the thermophile *Thermobifida fusca* (*T. fusca*) produces a full suite of enzymes required for lignocellulosic biomass processing.^23^ Of particular interest is the endocellulase Cel5A which has been previously engineered by our team for improved catalytic activity, and the focus of the work herein, the exocellulase Cel6B. Cel6B is a GH family-6 processive cellobiohydrolase that contains a native family-2a carbohydrate binding module (CBM).^22,24^ Processive exocellulases like Cel6B have a more nuanced mechanism that can be divided into initial binding to the bulk substrate interface, threading of the non-reducing cellulose substrate chain end into the active site and formation of the enzyme-substrate complex which commits the enzyme to processive motility cycles with cellobiose release.^25^ The full length CBM2a – Cel6B enzyme does have some shortcomings that limit its catalytic efficiency. For one, at the exit of Cel6B’s active site tunnel is a flexible loop that needs to open for expelling the cellobiose product that often leads to product inhibition.^26^ Single molecule tracking of CBM2a – Cel6B on cellulose also revealed that while the CBM is essential in assisting motility, the majority of the enzyme’s processive cycle is stalled on cellulose likely due to non-productive CBM binding where the stationary bound enzyme is more prone to denaturation.^27^ This effect has been observed with other cellobiohydrolases where prolonged substrate association leads to poor long-term stability and denaturation.^28^

There have been several previous efforts to engineer Cel6B and other GH6 exocellulases to improve functionality for cellulose degradation. Previous work with GH6 enzymes have targeted improving thermostability and resulting exo-exo synergism that resulted in a 2-fold increase in hydrolysis yield through SCHEMA^29^ recombination of several thermostable GH6 blocks.^30–32^ Targeting *T. fusca* Cel6B specifically, Vuong et al. targeted two mutations in the active site tunnel to improve processivity in an effort to increase synergism with endocellulase Cel5A.^33^ While they found the mutations beneficial for increasing processive run length, it did not successfully increase synergism with its endocellulase partner. One potential promising route for improving such exocellulases is through protein supercharging. Supercharging involves mutating several solvent exposed amino acid residues to negatively or positively charged residues to generate enzymes with high theoretical net charge,^34^ and has been successful in mitigating non-productive binding to lignin.^35,36^ This approach has been successfully deployed on endocellulase Cel5A and has demonstrated success in improving productive cellulose binding, catalytic activity, and thermostability for supercharged endocellulases.^37^ However, this method has yet to be utilized to engineer more active exocellulases.

Here we build upon our previous work with supercharging endocellulases and extend this computational protein design workflow to rationally design a supercharged library of Cel6B catalytic domains (CD) and its native CBM2a from *T. fusca*. We hypothesize that supercharging can improve the activity and stability of CBM2a – Cel6B and enhance endo – exo synergism by either improving individual hydrolytic activity of each enzyme partner or facilitating charge-guided colocalization of enzymes on the cellulose surface. To test this hypothesis, we designed and constructed a library of supercharged CBM2a and Cel6B CD constructs using Rosetta macromolecular software. A total library of 32 constructs including the native wild-type (WT) CBM2a – Cel6B fusion constructs were designed and synthesized spanning a full net charge range of −84 to +2 (pH 7.0) and activity of the entire library of mutants was assessed using a soluble cell lysate activity screen on several model soluble and insoluble cellulosic substrates. One key construct containing a supercharged CBM and the native CD, called D5 CBM2a – WT Cel6B, showed more than 2-fold increased activity on all substrates tested at pH 5.5 in our screen. Purified enzyme assays reveal this enzyme’s activity can be pushed to 2.3-fold higher specific activity compared to native enzyme on crystalline cellulose in presence of NaCl added, as well as no discernable changes in the optimal hydrolysis pH, a departure from what was observed with supercharged endocellulases in our previous study.^37^ These trends point to the fact that improved productive binding affinity is likely not the key driver for *T. fusca* exocellulase activity improvement. Thermal shift assays elucidate that, as a result of supercharging, for this particular construct, decoupling of the CBM and Cel6B CD melting points reveals a higher intrinsic melting temperature by nearly 18 °C. Furthermore, synergism with supercharged Cel5A endocellulases identified in our previous work showed that for both classes of enzyme, exocellulase and endocellulase, the constructs showing the highest activities individually were the best partners for synergistic cellulose hydrolysis.

## EXPERIMENTAL SECTION

### Reagents

The lignocellulosic biomass substrate primarily used in this study was AFEX pretreated corn stover prepared and provided by Dr. Rebecca Ong’s lab (Michigan Technological University) based on published protocols.^11^ Avicel (PH-101, Sigma-Aldrich) was utilized as a control for crystalline cellulose-I, as well for the preparation of phosphoric acid swollen cellulose (PASC) made in house based on established protocols.^38^ Genes used for expression of all wild-type and engineered constructions were synthesized and provided by the Department of Energy Joint Genome Institute (DOE-JGI). All other reagents used were procured from either Sigma Aldrich or Fisher Scientific unless otherwise noted in subsequent sections.

### Computational design of CBM2a – Cel6B library and plasmid generation

Although belonging to the same CBM family (2a) and sharing significant homology, the CBM2a construct described in this work is distinct from the one designed in our earlier Cel5A study.^37^ As a solved crystal structure was not available for the native Cel6B CBM2a used here, a homology model was constructed using Rosetta CM.^39^ The structure of the CD for *Thermobifida fusca* Cel6B has been solved (PDB: 4B4H)^24^ and was used for generating all supercharged Cel6B CD designs. Protein supercharging was completed following the same methodology described in our previous work using Rosetta Macromolecular modelling software.^37^ In short, mutations to either positive (K and R) or negative (D and E) amino acid residues were iteratively made to surface exposed amino acid residues on both the CBM and Cel6B catalytic domain individually to generate constructs with high theoretical net negative or positive charge. Both AvNAPSA (Average number of Neighboring Atoms Per Side-chain Atom)^40^ and Rosetta supercharging^41^ protocols were utilized to define the protein surface and introduce mutations. The motivation in including both methodologies was to avoid biasing designs based on design principle and computational protocol. Additional user defined stipulations were included within a resfile to avoid introducing deleterious mutations by preserving residues; (i) within 10 Å of the CBM binding face, (ii) within 10 Å of the Cel6B CD active site, (iii) that possess the correct charge base, (iv) that cap alpha helices, and (v) form salt bridges. Simulations were run until a target net charge was reached, and output structures were analyzed in PyMOL to ensure the flags set in the resfile were not violated. Nucleotide sequences for the final designs were codon optimized for *E. coli* expression and provided to the Joint Genome Institute (Department of Energy) for gene synthesis. Synthesized sequences were cloned between the KpnI and XhoI restriction sites of the pET45b(+) expression vector (www.addgene.org/vector-database/2607/) and transformed into the T7 SHuffle (New England Biolabs) expression strain. Each construct was individually received as a 20% glycerol stock ready for expression and characterization as described in the subsequent sections.

### Expression for cell lysate screening

A total of 32 CBM2a – Cel6B constructs including the wild-type enzyme were received and grown as 200 mL expression cultures for initial characterization of the entire library. Glycerol stocks were swiped and used to inoculate 10 mL of LB media and 100 µg/mL carbenicillin and grown at 37 °C with 200 RPM orbital shaking for 16 hours. A larger bank of glycerol stocks was prepared from these overnight cultures to avoid repetitive thawing of the source plate. The remaining inoculum was transferred to 200 mL of Studier’s auto-induction medium (TB+G)^42^ with an additional 100 µg/mL carbenicillin. Cultures were incubated for an additional 6 hours at 37 °C to provide sufficient time for cells to consume glycerol and glucose present in the media and reach exponential growth before inducing expression in two steps: first 25 °C for 24 hours followed by 16 °C for 20 hours. Cells were isolated from spent media via centrifugation at 30,100 x g for 10 mins at 4 °C in a Beckman Coulter centrifuge. Dry cell pellets were kept in 50 mL falcon tubes and stored at −80 °C until used for characterization.

### Preparation and characterization of soluble cell lysates

Soluble cell lysates were prepared for screening library activity by first harvesting two separate 0.5g aliquots of cells from the main dry pellet for the entire library. Each aliquot was resuspended in either lysis buffer A (50 mM sodium acetate buffer, 10 mM NaCl, and 20% (v/v) glycerol, pH 5.5) or buffer B (50 mM phosphate buffer, 10 mM NaCl, and 20% (v/v) glycerol, pH 7.5), 35 µL protease inhibitor cocktail (1 µM E-64, Sigma Aldrich), and 2.5 µL lysozyme (Sigma Aldrich). Cells were lysed via sonication with a Qsonica Q700 sonicator with 1/8” microtip probe for 1 minute (Amplitude = 20, pulse on time: 5 s, pulse off time: 30 s) on ice to avoid overheating. Soluble cell lysates were clarified from insoluble cell debris by centrifugation at 15,500 x g for 45 minutes in an Eppendorf 5424 centrifuge with rotor FA-45-24-11.

The soluble cell lysates were used to screen for exocellulase activity of each construct in two different lysis buffers on AFEX pretreated corn stover, cellulose-I, and PASC. All insoluble substrates were prepared as slurries in deionized water (DI) water with 0.2 g/L sodium azide to prevent microbial contamination. AFEX corn stover was milled to 0.5mm and prepared as a 25 g/L stock, cellulose-I (Avicel PH-101) was prepared as a 100 g/L slurry, and PASC was prepared by swelling cellulose-I with phosphoric acid and precipitating it in ice-cold DI water to a final concentration of 10 g/L. Hydrolysis assays were performed by adding 100 µL of substrate slurry (AFEX, cellulose-I, or PASC) to 100 µL of crude cell lysate in a 0.2 mL 96-well round bottom microplate (Greiner Bio-One). No additional buffer was added as lysates were prepared in buffers with the desired pH to test activity. All conditions were tested in quadruplicate and reaction blanks were included replacing soluble cell lysates with the generic lysis mixture (lysis buffer, PIC, and lysozyme), and lysed T7 SHuffle with an empty vector was used as a negative control to ensure there was no background hydrolysis occurring from heterologously expressed proteins. Plates were sealed with a TPE capmat-96 (Micronic) plate seal and taped tightly with packing tape along edges to ensure no evaporative losses during incubation. Hydrolysis plates were incubated at 60 °C with 5 RPM end-over-end mixing in a VWR hybridization oven for six hours. This temperature was chosen based on prior work that found 60°C to be the optimal temperature for *T. fusca* cellulases^37,43^. An excess amount of substrate was used in these assays to allow for saturation with available enzyme activity sites and absolute activity to level off. The concentration of reducing sugars released by enzymes in the soluble hydrolysate was estimated by dinitrosalicylic acid (DNS) assay,^44^ as described in previous work,^37,43^ and compared to glucose standards.

### Scaled up expression and protein purification

The full length CBM2a-Cel6B construct exhibit several challenges in downstream purification relating to his-tagged truncation products as well as expressed heat shock chaperones.^45^ In order to produce sufficient amount of native enzyme (WT CBM2a – WT Cel6B) as well as the best performing supercharged constructs (e.g., D5 CBM2a – WT Cel6B), all relevant constructs were expressed on a 6.5L scale in an 10L, water chilled glass bioreactor controlled by an Eppendorf BioFlo 120 control tower. Both constructs were transformed into a BL21 (DE3) expression strain (New England Biolabs) as optimization testing found higher purified protein yields with this strain compared to the original T7 SHuffle. Briefly, 250 mL of sterile LB media with 100 µg/mL carbenicillin was inoculated with glycerol stock and incubated for 16 hrs at 37 °C. In parallel, 6.5 L of TB+G media^42^ was prepared within the glass vessel of the bioreactor. After calibrating both dissolved oxygen (DO) sensor and pH probe (Mettler Toledo), the vessel was sealed and any sampling tubes plugged and covered with aluminum foil. The entire reactor assembly and tubing used for base addition during incubation were sterilized by steam autoclave. The 250 mL overnight inoculum was added to the reactor with an additional 100 µg/mL carbenicillin and incubated for four hours at 37 °C with temperature maintained by a heating jacket until an exponential OD was reached. At this point, reactor temperature was decreased to 25 °C and protein expression was auto induced for 24 hours. For the duration of incubation, agitation rate and air sparing (compressed bench air) were controlled by DO cascade. A DO setpoint of 30% was maintained by manipulating agitation rate first from 300 RPM to 450 RPM, followed by air sparging which ranged from 0.1 SLPM to 2.5 SLPM. The starting pH of the media was 7.4, and a setpoint of pH 7.0 was maintained by addition of sterile 5% NaOH. A setpoint of 7.0 was chosen as to avoid excessive addition of base at the risk of killing the cell culture. Finally, to combat foaming, 30% Antifoam B (Sigma-Aldrich) was added incrementally as needed detected by a wet vs. dry condition of a liquid level sensor.

For the other constructs tested, especially those with two supercharged domains or domains with high theoretical net charge, standard shake flask cultures were used for expression in the original T7 SHuffle expression strain. For these constructs, 10 mL of LB media with 100 µg/mL of carbenicillin was inoculated with a frozen glycerol stock and incubated for 16 hours at 30 °C. The inoculum was added to 500 mL of sterile LB media with 100 µg/mL of carbenicillin and incubated for ∼4hr at 30 °C until the OD reached 0.4-0.6 at which point 200 µL 1M (Isopropyl β-d-1-thiogalactopyranoside) IPTG was added for an effective concentration of 0.4 mM IPTG to induce expression at 16 °C for 24 hours. After expression, for both either reactor or shake flask cultures, cell pellets were harvested via centrifugation at 30,100 x g for 10 mins at 4 °C in a Beckman Coulter centrifuge. Cell pellets were resuspended in 15 mL of cell lysis buffer (20 mM phosphate buffer, 500 mM NaCl, and 20% (v/v) glycerol, pH 7.4), 200 µL PIC, and 15 µL lysozyme for every 3 grams of dry cell pellet. Cells were adequately resuspended by vortexing until homogenous and allowed to incubate on ice for 30 mins to chill the lysate mixture and allow time for the lysozyme present to begin breaking down the cell membrane. Cells were lysed via sonication on ice with a Qsonica Q700 sonicator with a 1/4” microtip for 5 minutes (Amplitude = 10, pulse on time: 10s, pulse off time: 30s). After sonication, the soluble cell lysate was isolated from cellular debris via centrifugation in an Eppendorf 5810R centrifuge at 12,000 x g for 45 mins at 4 °C.

A two-step purification process was employed for isolating his-tagged CBM2a-Cel6B enzymes from background *E. coli* proteins. The first step of purification used was immobilized metal affinity chromatography (IMAC) using a BioRad NGC FPLC with a His-trap HP Ni^2+^ – NTA column (Cytiva). System plumbing was cleaned with 1 M NaOH prior to loading any sample followed by rigorous washing with DI water and equilibration with buffer A (100 mM MOPS, 500 mM NaCl, 10 mM imidazole, pH 7.4). Lysate samples contained no imidazole, thus enough buffer B (100 mM MOPS, 500 mM NaCl, 500 mM imidazole, pH 7.4) was added so that the effective imidazole concentration before column binding was 20 mM. Cell lysates were loaded on the column at a rate of 1 mL/min and column flow through was collected and analyzed via SDS-PAGE to ensure there was minimal protein loss during binding. Upon loading sample, the column was washed with 10 – 20 column volumes of buffer A until a stable baseline was achieved. Proteins were eluted in three distinct isocratic steps with increasing concentration of buffer B: (i) 5% buffer B, (ii) 10% buffer B, and (iii) 100% buffer B and all eluent fractions were collected corresponding to A280 peaks and analyzed for SDS-PAGE. Purity of the final full-length CBM2a-Cel6B construct in the 100% buffer B eluent is only roughly 60% based on gel image densitometric analysis, with the primary contaminant being a his-tag containing truncation product. Attempts to further separate this product from the full-length enzyme using ion exchange chromatography were unsuccessful, likely due to similar pIs for the full-length enzyme and smaller truncation product.

Due to the disparity in molecular weights between the full-length enzyme and truncation product, size exclusion chromatography (SEC) was employed as a second step of purification using a HiPrep Sephacryl S-200 HR preparative SEC column (Cytiva) with 26 mm inner diameter. Prior to sample purification, the column was cleaned with one column volume (320 mL)1M NaOH at 1.3 mL/min and washed with 5 column volumes of DI water at 2.6 mL/min on a BioRad NGC FPLC. IMAC eluents for each construct were pooled and concentrated to below 13 mL using a Vivaspin 20 10kDa cutoff centrifugal concentrator (Cytiva). The SEC column was equilibrated with two column volumes of 50 mM MOPS pH 7.5 with 100 mM NaCl and sample was loaded at 1.3 mL/min. Protein samples were eluted by equilibrating the column with one column volume of the same SEC buffer at 1.3 mL/min with constant eluent collection in 5 mL aliquots. Fractions corresponding to A280 peaks were saved and analyzed with SDS-PAGE to identify the aliquots containing >90% purity CBM2a-Cel6B. These samples were pooled and concentrated 5-fold in a Vivaspin 20 10kDa cutoff centrifugal concentrator (Cytiva) before being aliquoted in 100 µL fractions, flash frozen in liquid nitrogen, and stored at −80 °C for further characterization.

### Purified enzyme hydrolysis assays

Catalytic activity of the native CBM2a-Cel6B enzyme and selected supercharged constructs was assessed on AFEX corn stover, crystalline cellulose-I, and PASC testing responses in activity to different solution pH as well as ionic strength tolerance. Although processive exocellulases like Cel6B are often subject to cellobiose product inhibition,^46–48^ the final cellobiose product yield in our assay configuration was low enough as to not significantly inhibit Cel6B based on preliminary tests. Therefore, a glucosidase enzyme was not included in these assays since its activity would also be altered by the different solution conditions tested herein. All reactions were conducted in 0.2 mL round bottom microplates (Greiner Bio-one) at a constant enzyme loading of 120 nmol enzyme per gram of substrate chosen based on previous research with *T. fusca* cellulases^37^ including Cel6B.^43^ Reactions with AFEX corn stover used 2 mg total of substrate, cellulose – I reactions contained 4 mg of substrate as preliminary assays showed low activity due to limited cellulose chain end availability for the exocellulase to initiate hydrolysis on. Assays sweeping solution pH contained 10 µL of 1 M stock buffer (either sodium acetate or sodium phosphate ranging from pH 4.5 – 7.5) for an effective buffer concentration of 50 mM, 50 µL of cellulase dilution (120 nmol/g loading), and the volume was made up to 200 µL with DI water. Hydrolysis plates were sealed with a TPE capmat-96 (Micronic) plate seal and taped tightly with packing tape to ensure a tight seal along the edges of the plate seal mat. Plates were incubated for 24 hours at 60 °C in a VWR hybridization oven with 5 RPM end-over-end mixing. Assays testing the effect of salts on activity contained the same amount of substrate and enzyme. These assays were tested in 50 mM potassium phosphate, pH 6.0 which was the identified optimum for all of the purified enzymes, with the addition of 100 mM NaCl (effective concentration) in reaction wells. Volume was kept constant at 200 µL with DI water, and plates were sealed and incubated with the same conditions described for the pH sweep. After incubation, microplates were centrifuged to settle all insoluble solids and reducing sugar equivalents in soluble hydrolyzates were estimated by DNS assay in the same manner described in the previous sections. All conditions were tested in quadruplicates and water/buffer blanks were used to subtract background absorbance from the final DNS response.

### Thermal shift assay and data processing

Thermal shift assays were conducted based on manufacturer protocols in a Bio-Rad CFX Opus 96 Real-time PCR system. Briefly, 5 µL of 50x SYPRO orange dye (Thermo Fischer Scientific), 2.5 µL of 1 M sodium phosphate buffer pH 6.0, 17.5 µL of DI water, and 25 µL of 10 µM protein dilution were added to a clear low-profile Hard-Shell 96-well PCR plate (Bio-Rad) and sealed with an optically clear adhesive Microseal ‘B’ plate seal (Bio-Rad). System software was configured based on Bio-rad’s provided protein thermal shift protocol, and thermostability was accessed from 25 – 99 °C with a 0.5 °C increment and 5s read time. Each enzyme was tested in quadruplicate, and a buffer blank was included to subtract background fluorescence from protein melt curves. Data was analyzed outside of the Bio-Rad software in MATLAB. Replicates were averaged and background fluorescent signal from the blank was subtracted at each temperature tested. For more accurate melting temperature estimation, cubic spline interpolation was used on the dataset to smooth the melt curves and interpolate data at 0.001 °C intervals. Melting temperatures were estimated by taking the negative first derivative of the interpolated data set using the MATLAB gradient function and melting points were found using MATLAB’s min function.

### Exo-endo cellulases synergism assays

Synergism between endo and exo cellulolytic enzymes was assessed by testing the activity of the native CBM2a – Cel6B and most active supercharged construct (D5 CBM2a – WT Cel6B) with the native and supercharged CBM2a – Cel5A constructs identified from our previous work.^37^ As this minimal cellulase mixture is expected to synergize and produce a higher hydrolysate yield, a β-glucosidase (Bglc) from *T. fusca* (genes also synthesized and provided by JGI) was expressed and purified in a manner similar to described here and included in assays at 10% of the total cellulase loading. AFEX pretreated corn stover and crystalline cellulose – I substrates were both used as substrates with a total of 2 mg of substrate present in reaction wells. To assess synergistic improvements of the endo-exo mixture, individual enzyme activity was first screened at 120 nmol/g with the glucosidase present. Synergism was assessed by modifying the amount of cellulase present in reaction wells testing a 1:3, 1:1, and 3:1 endo:exo cellulase molar ratio while keeping the total enzyme loading constant at 120 nmol enzyme per gram substrate (along with 12 nmol glucosidase/g substrate). Reaction plates were caped with a TPE capmat-96 (Micronic) plate seal and taped tightly to prevent evaporation before incubating 24 hours at 60 °C with 5 RPM end-over-end mixing in a VWR hybridization oven. The same buffer composition as was used in the salt screen (50 mM phosphate pH 6.0) was used for all assays and experiments were repeated in quadruplicates. Reducing sugar equivalents in the soluble hydrolyzate were estimated via DNS assay following the same protocols mentioned earlier.

## RESULTS AND DISCUSSION

### Computational design of supercharged library

The native CBM2a – Cel6B enzyme possesses a net charge of −4 on the CBM and −20 on the Cel6B catalytic domain based on the count of positive and negative amino acid residues present in the primary amino acid sequence. As Cel6B natively possesses a high dispersion of negative charged residues, several mutations were required to produce constructs with a positive charge while extremes of positive charge were near impossible to reach without risking significant destabilization of the catalytic domain. As stated in the methodology section, supercharging was completed using both AvNAPSA and Rosetta Supercharging (Rsc) protocols in order to produce sequence diversity between supercharged constructs since both methods will predict different mutations. AvNAPSA is much more conservative in scoring residues for mutations, with only polar amidic and charged residues being mutated.^40,41^ For this reason, AvNAPSA produced designs with only modest theoretical net charges. AvNAPSA designs also included some degree of mutation redundancy where constructs with higher theoretical charge included mutations from the previous design with lower net charge. Rosetta supercharging allowed for larger sampling of the design space with designs at charge extrema being produced using this method.

A total of eight supercharged CBM2a constructs were designed spanning a net charge range of −13 to 4 with an equal dispersion of negative to positive designs (**Figure 1**). Of the total eight constructs, constructs D3 – D6 were designed with AvNAPSA and constructs D1, D2, D7, and D8 designed with Rsc. AvNAPSA designs possess modest charges and redundancy where design D3 includes the same mutations as D4. All mutations made for these constructs were to polar amidic residues to either aspartic or glutamic acid based on residue scoring in Rosetta software. Rsc designs included mutations to surface exposed serine and threonine residues as well as polar amidic residues to access higher charge ranges. Since two methods were used, the goal in targeting net charge ranges was to include enough granularity to sample a wide charge range, but not to span large differences in net charges when switching between methods to ensure differences in resulting activity are more biased to surface charge and not design methodology. This same principle was adhered to in our previous work producing results highly correlated to overall net charge and not biased to AvNAPSA or Rsc protocols.^37^ For this reason, constructs D2 and D3, and D6 and D7 are similar in overall charge.

**Figure 1.**
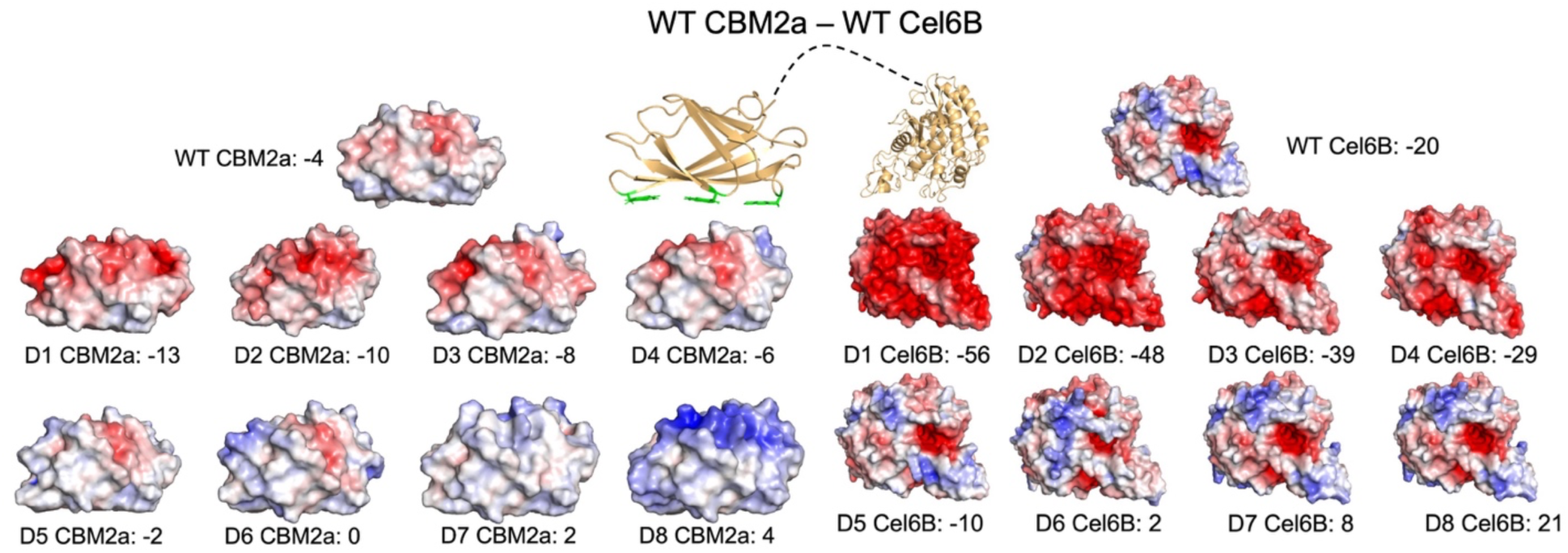
Computational design and library construction of CBM2a – Cel6B Library. Rosetta macromolecular software was used to identify solvent exposed amino acid residues on the surface of both CBM2a and Cel6B for mutation to positively charged (K, R) or negatively charged (D, E) amino acids. Each domain was mutated individually then one of eight CBM designs was fused with one of eight Cel6B CD designs with the same flexible linker peptide native to the full-length enzyme. Protein structures were generated in Alpha Fold using primary amino acid sequences, and output structures were relaxed using Rosetta’s fast relax function. Net charges for each design or wildtype domain (WT) are indicated in the figure and were calculated by counting the total number of charged amino acid residues. Electrostatic potential maps ranging from −5 kT/e (red) to +5 kT/e (blue) were generated using Adaptive Poisson – Boltzman Solver (APBS) plugin in PyMOL.

As the Cel6B catalytic domain (420 residues) is nearly four times larger compared to the CBM (109 residues), the potential design space allows for sampling at much higher charge extremes. Once again, eight total constructs were designed for the CD spanning a net charge range of −56 to +21 with an equal dispersion of AvNAPSA and Rsc designs. Mutations were made to the CD in a similar fashion as was described for the CBM, with majority of the positive mutations being made to lysine based on Rosetta scoring. Additionally, CD designs produced with Rsc at high charge extremes specifically targeted mutations to charged residues as this effectively increases net charge by ± 2 as compared to mutations made to uncharged residues. Including the wild type CBM and CD, there was a final total of 9 CBM and 9 CD constructs generated. The mutations made to generate each CBM and Cel6B CD construct are summarized in **Supplementary Table T1** and **T2** respectively. To construct the full-length CBM2a – Cel6B enzyme, a CBM construct would be fused to a Cel6B construct with the same flexible linker peptide native to the wild-type CBM2a – Cel6B enzyme. Of the 81 possible combinations (including native), all single domain mutants possessing one supercharged domain (CBM or CD) were synthesized, along with a range of combinatorial mutants to cover the design space omitting those with redundant net charges. The final library size of 32 constructs was synthesized by the JGI and received as glycerol stocks with recombinant DNA inserted into the pET45b(+) vector. To ensure errors were not made in synthesis or sample handling, plasmids for the library were extracted and sequenced prior to proceeding with expression and characterization. Nucleotide sequences for the synthesized library as well as corresponding amino acid sequences are provided in the sequences supplemental excel file.

### Comprehensive library screening based on soluble cell lysate activity

The entire supercharged Cel6B library was first expressed on a small scale and activity screened from soluble cell lysates to identify constructs with better catalytic performance compared to the native enzyme. This approach was previously successful with endocellulase Cel5A^37^ with two key limitations: (i) the pH chosen in this study underrepresented activity for some mutants, and (ii) the high salt concentration screened some supercharged interactions altering the activity for some constructs. Therefore, a modified approach was taken in this study. First, preliminary screening was completed with both the wildtype Cel6B enzyme and some supercharged Cel5A constructs from the previous study with altered lysis buffer concentrations. The key findings from these assays was that glycerol in the buffer did not produce adverse effects in activity, selecting two different buffer pH (5.5 and 7.5) provides a better representation of the activity for the broad library of constructs, the concentration of NaCl can be lowered to 10 mM to ensure solution stability without impacting activity, and increasing substrate load and incubation time will allow for absolute activity to plateau. Initially, it was desired to report activity as specific activity, but this is highly reliant on a defined enzyme concentration which is difficult to estimate from crude cell lysates. Although purification of cell lysate samples showed no significant differences in expression yields, increasing substrate concentration and incubation time is an added fail safe to ensure slight expression differences do not provide significant differences in the output activity.

The activity of the entire supercharged library on AFEX pretreated corn stover at pH 5.5 is depicted in **Figure 2A** and activity at pH 7.5 in **Figure 2D**. The initial observation from this dataset is that supercharging generally produced constructs with similar activity to the wild-type enzyme (construct 1), with some displaying significantly higher hydrolysis yields. Those constructs displaying significantly lower activity on biomass all were negatively supercharged, likely due to poorer initial binding commitment to cellulose. Prior studies with supercharged CAZymes has shown a tradeoff when analyzing biomass activity; negative supercharging is beneficial to prevent binding to the negative structural polymer lignin, but to an extreme will also prevent adhesion to cellulose, ultimately preventing effective catalysis.^35–37^ This same effect is likely what is observed here as well for several negatively supercharged constructs. Initial observations point to positive supercharging being more beneficial which was also observed with Cel5A.^37^ At pH 5.5, the D5 CBM2a – WT Cel6B construct was the most promising (construct 13). This construct contains a slightly positive supercharged CBM fused to the native Cel6B enzyme displayed a 1.7-fold improvement in activity compared to the native enzyme. This was closely followed by D5 CBM2a – D5 Cel6B (construct 22) which combines the same D5 CBM2a with a positively supercharged D5 Cel6B. As was seen in the past when combining two supercharged domains, the overall benefits of the individual supercharged domains is not additive, but instead, the activity is capped by whichever individual domain showed the highest activity.^37^ This again appears to be the case for D5 CBM2a – D5 Cel6B where the activity closely resembles that of the more active performing D5 CBM, with a slight drop in activity likely due to the poorer performance of the D5 Cel6B domain (construct 5). The motivation for testing activity at pH 5.5 is that it is the reported pH optimum for many *T. fusca* cellulases like Cel6B and should give a closer translation to purified enzyme activity in ideal conditions. However, enzymes will experience a sharp pH gradient when lysing *E. coli* cells with a cytoplasmic pH of 7.4 – 7.9^49^ into pH 5.5 buffer which may compromise stability for several constructs. Therefore, a second buffer at pH 7.5 was used in order to provide a more stable solution condition during lysis at the potential expense of activity. D5 CBM2a – WT Cel6B does display this decrease in activity at pH 7.5 more closely resembling the activity of the native enzyme. At this pH, the effects of positive supercharging are effectively nullified as the binding module become more negatively charged in solution, an effect also observed previously with other supercharged enzymes.^37^ It is also important to note that the background proteins present in the cell lysate may have a stabilizing effect as they interact with recombinant Cel6B constructs. This can be observed for constructs that have high theoretical charge which appear more active at pH 7.5 where they showed much poorer catalysis at pH 5.5. There does not appear to be any trend relating charge to the output activity for these constructs, however.

**Figure 2.**
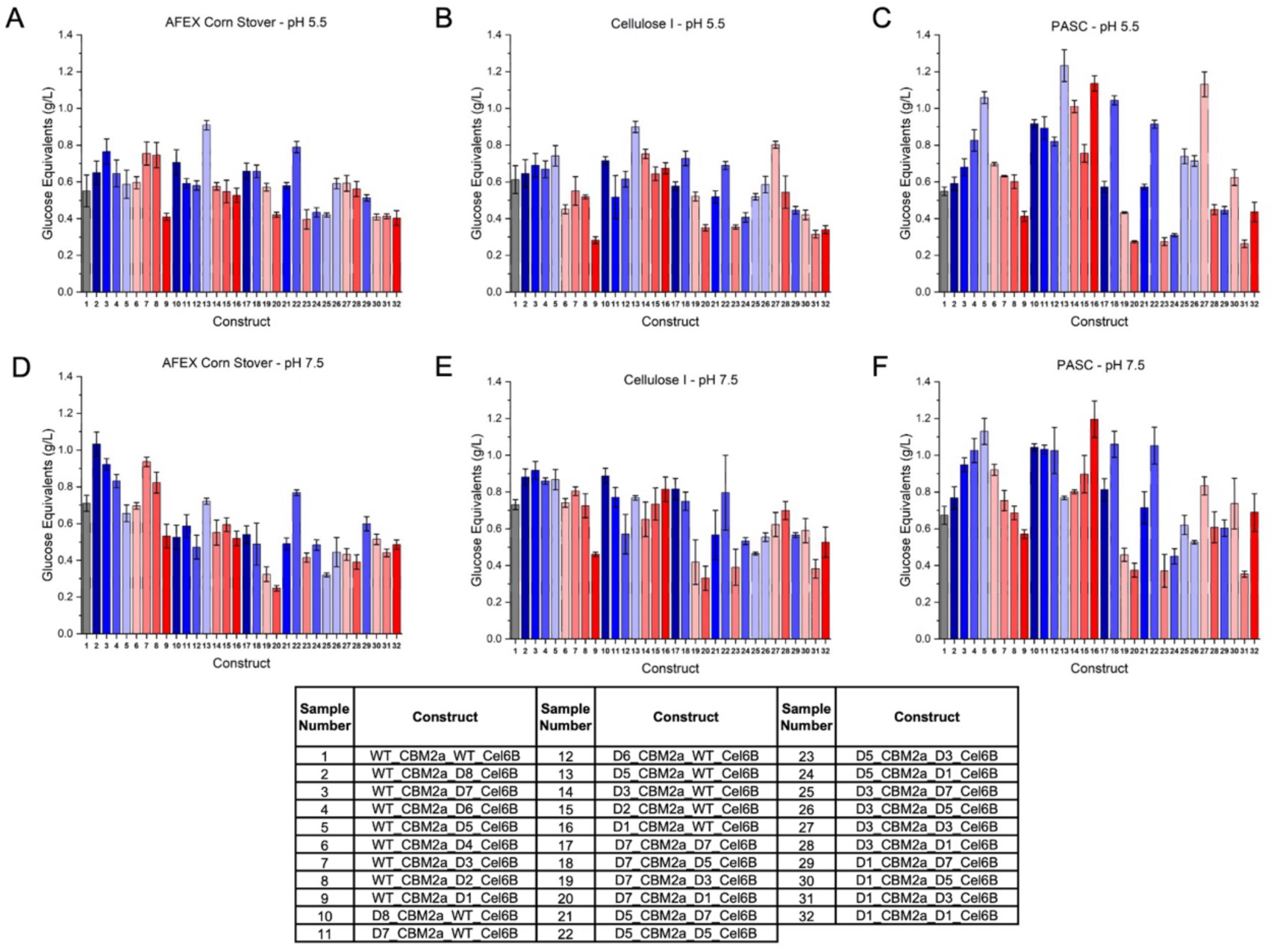
Screening of supercharged Cel6B constructs from soluble cell lysate reveals improved hydrolytic performance on biomass and model cellulose substrates. The entire CBM2a – Cel6B library was expressed as 200 mL auto-induction cultures and 0.5g aliquots of the resulting dry cell pellet was sonicated in buffer containing either (A-C) 20 mM sodium acetate pH 5.5 or (D-F) 20 mM sodium phosphate pH 7.5 buffer with 10 mM sodium chloride and 20% (v/v) glycerol. Soluble cell lysates were used to screen activity on (A, D) Ammonia fiber expansion (AFEX) pretreated corn stover, (B, E) crystalline cellulose – I (Avicel PH-101), and (C, F) phosphoric acid swollen cellulose (PASC). AFEX corn stover was prepared as a 25 g/L slurry, cellulose-I was prepared as a 100 g/L slurry from Avicel PH-101, and PASC was prepared as a 10 g/L slurry all in DI water. Activity was tested on all substrates by incubating 100 µL of substrate slurry with 100 µL of cell lysate for 6 hours at 60 °C with 5 RPM end-over-end mixing. Reducing sugar equivalents were estimated using DNS assay and compared to glucose standards. All data points represent the average of four technical replicates and error bars represent one standard deviation from the mean. Constructs are color coded based on net charge with most negative in dark red and most positive in dark blue.

Hydrolysis of crystalline cellulose – I (**Figure 2B, E**) provides insight into the interactions solely with cellulose as opposed to biomass where hemicellulose and lignin are also present. Once again, at pH 5.5, D5 CBM2a – WT Cel6B exhibits the highest activity with a 1.5-fold improvement compared to the native enzyme. Remarkably, these improvements are being observed at relatively high solids loading (50 mg/mL) where CBMs have been thought to be less useful due to the abundance of substrate.^50,51^ Analyzing activity for the constructs containing only a supercharged CD (2-9), those positively supercharged show significantly better performance compared to their negatively supercharged counterparts. This is to be expected due to the slight negative surface charge found on cellulose^36^ which the negative CD’s repel. Still, none show much improvement compared to the native enzyme. Comparing constructs with only a supercharged CBM (10-16), we see less detriment with the negatively charged constructs. This is likely due to the aforementioned high solid loading environment where the detrimental impact of cellulose repulsion on only a binding domain is less pronounced. For these constructs, the native Cel6B CD has ample substrate to bind to even with unfavorable interactions on the assisting binding domain. Interestingly, the positively supercharged CBMs showed poorer activity with exception to the best performing construct D5 CBM2a – WT Cel6B. This is likely due to the impact of too high of a binding affinity on cellulose. As Cel6B processively degrades cellulose, previous work has shown that the CBM assists in this processive motion.^27^ Higher affinity CBMs may begin to oppose this processive action by remaining too tightly bound to cellulose or binding tightly in non-catalytically relevant conformations; an effect that has been observed in other processive exocellulases like Cel7A.^52^ This also suggests that the activity effects observed for D5 CBM2a – Cel6B may not solely stem from improved cellulose binding as was the case with supercharged Cel5A constructs^37^ but instead by some intrinsic stabilization occurring as a byproduct of mutations introduced. At pH 7.5, these trends generally remain the same, except differences in activity compared to the native enzyme are more subtle, likely due to the same stabilization with the background lysate discussed with biomass. A full statistical analysis of supercharged construct activity compared to the native enzyme is provided in the supplemental information and is viewable in **Supplementary Figure S1.**

Activity on PASC is quite interesting because it presents different catalytic opportunities compared to cellulose-I and AFEX corn stover. PASC is a much more endoglucanase friendly substrate that is significantly amorphous, and easily hydrolysable without the need for enzyme pairs or binding domains. For this reason, PASC was used as a positive control to ensure C-terminal catalytic domains were being expressed in the event that activity was not observed on biomass and crystalline cellulose. Still, there are some intriguing trends to note. Again, at pH 5.5, D5 CBM2a – WT Cel6B exhibits the highest overall activity, and for several constructs at both pH 5.5 and 7.5, there are 2-fold and greater improvements in activity. This suggests that some supercharged constructs may be more endo-active than the native enzyme. Additionally, when analyzing the activity trends for constructs 2-9, there is clearly a net charge preference where activity follows a unimodal distribution indicative of a Sabatier optimum.^53–55^ This again suggests a great endo active contribution from these enzymes as this same trend was readily apparent when supercharging endocellulase Cel5A.^37^ Interestingly, the peak of this volcano plot falls on the WT CBM2a – D5 Cel6B construct. This seems to suggest that the performance observed on biomass and crystalline cellulose for D5 CBM2a – D5 Cel6B may be a trade-off from the beneficial stabilization improving exo-activity with the D5 CBM and the improved endo activity for D5 Cel6B, especially on pretreated biomass where amorphous regions are accessible. When comparing the overall trends when supercharging only CBM, CD, and combining them (**Supplemental Figure S2, S3**) the greatest difference in activity occurs when supercharging only the CBM. These constructs showed a larger disparity in activity compared to supercharging only the CD. In nearly every case observed, combining oppositely supercharged domains significantly hinders catalytic performance, likely due to the two domains interacting with one another at the cost of substrate engagement, which was also observed in previous studies.^37^

### D5 CBM2a – WT Cel6B shows more than 2-fold improvements in activity in purified enzyme assays

The cell lysate screen provided a rough estimate of activity of the entire library with the insight that construct 13, D5 CBM2a – WT Cel6B standing out as the clear winner within the library. This construct, as well as several others that showed higher activity on at least two substrates were expressed on a large scale and purified for closer characterization. Initially, assays sweeping different buffer pH were ran in order to analyze the effect that changing pH has on Cel6B. Results (**Supplemental Figure S4**) showed no shift in the optimal hydrolysis pH for any of the supercharged constructs compared to the native enzyme. In fact, on AFEX pretreated corn stover, there is a steep drop off in activity outside the pH 5.5 – 6.0 range. This is a stark contrast to what was observed when supercharging an endocellulase like Cel5A. For Cel5A, slight differences in surface charge shifted the pH optimum of hydrolysis for each supercharged construct.^37^ This trend does not hold for Cel6B. It appears that the initial binding commitment to the bulk substrate is not rate limiting in the case here,^25^ and thus, Cel6B is likely not improved through finding a Sabatier optimum^54^ with an intermediate cellulose binding strength like Cel5A was (**Supplemental Figure S5**).

In order to assess if the supercharged constructs behave differently in the presence of salts, hydrolysis assays with AFEX and cellulose-I were run both with and without 100 mM NaCl present. A pH of 6.0 was once against used here based on the results from pH sweeps. When hydrolyzing AFEX corn stover (**Figure 3A**) the most striking observation is the overall low activity for all enzymes. Hydrolysis yield did not exceed 0.1 g/L, unlike in the lysate screen where reducing sugar yields approached 1 g/L for the D5 CBM2a – WT Cel6B mutant. This difference is likely related to the presence of background proteins in the lysate helping to block both lignin and cellulose helping to alleviate non-productive binding for these cellulases in lysate assays.^20^ For the purified enzymes, no such sacrificial proteins are added, and thus they will be further limited by this nonproductive binding. For the two highly positive constructs (D7 CBM2a – WT Cel6B and D7 CBM2a – D5 Cel6B), a much lower activity is observed compared to the native enzyme, likely due to high adsorption to lignin, or too high of a propensity for binding cellulose. The negatively charged mutant D3 CBM2a – WT Cel6B has the inverse problem, poorer cellulose adhesion due to the negative charge. The chosen winner from the lysate screen, D5 CBM2a – WT Cel6B shows a slightly lower activity compared to the native enzyme without salt, and this is rescued with salt present, likely screening electrostatics interactions with lignin. Still, there is no significant improvement compared to the wild-type enzyme.

**Figure 3.**
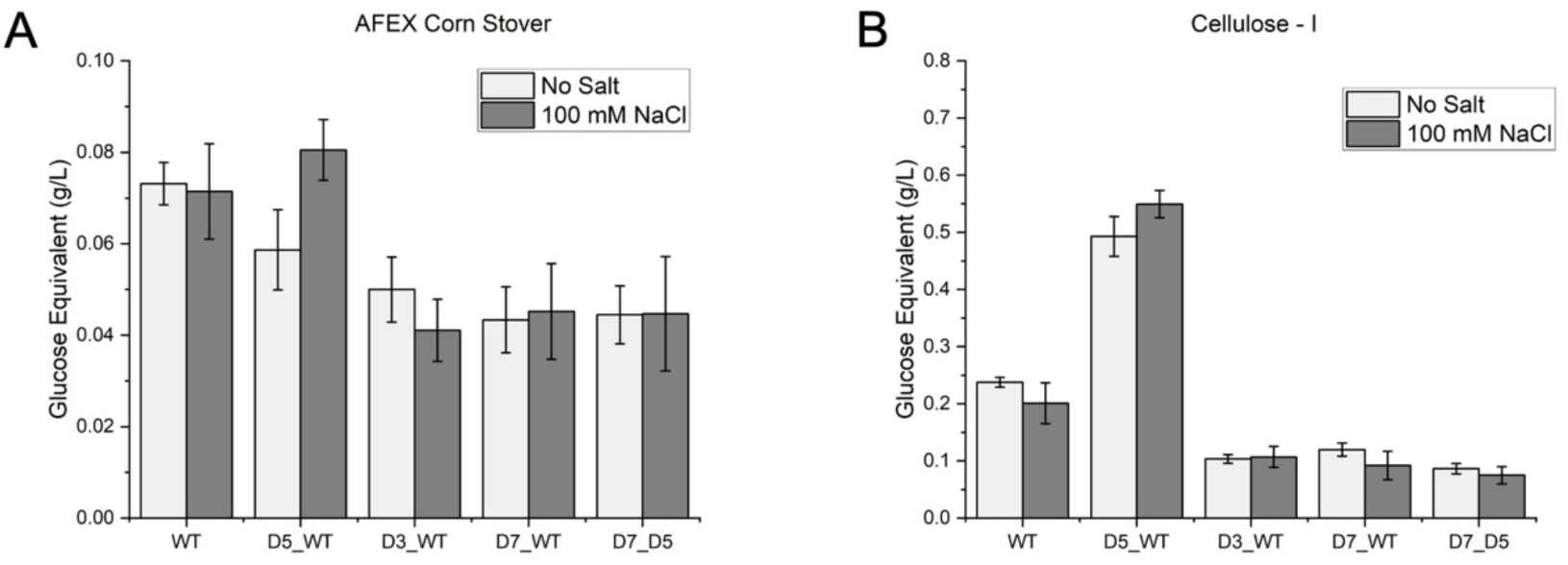
Moderate positive supercharging enhances Cel6B activity under ionic conditions, while extreme charge alters enzyme–substrate interactions. Enzymes were expressed as auto-induction cultures in a construct dependent manner as described in the experimental section. N-terminus his-tagged enzymes were purified from *E. coli* lysate in two steps first by immobilized metal affinity chromatography then by size exclusion chromatography. Assays were conducted both with and without 100 mM sodium chloride present on (A) AFEX corn stover and (B) crystalline cellulose – I. All enzyme dilutions were made in DI water and a constant enzyme-substrate loading of 120 nmol / g substrate was used for assaying activity. Hydrolysis was conducted at 60 °C for 24 hours with 5 RPM end-over-end mixing for 24 hours and reducing sugar equivalents were estimated via DNS assay and compared to glucose standards. Enzyme construct is denoted on the x-axis with the native full-length CBM2a – Cel6B labeled WT and supercharged constructs annotated as CBM design number_ Cel6B design number (or native/WT). All data reported represents the average of four technical replicates, and error bars represent standard deviation from the mean.

Activity assay results on cellulose-I show a different outcome for this enzyme however (**Figure 3B**). Without salt, a clear improvement in activity is observed compared to the native enzyme. This does not appear to be driven by initial binding because this activity increases with the addition of salt with an overall 2.3-fold increase in activity compared to wild-type. If this improvement was due to favorable binding between the positive charged CBM and negative cellulose surface, the addition of salt would screen this effect and a decrease in activity would be observed. Instead, it is likely that some intrinsic stability as a result of supercharging is leading to the improvements for this construct. Previous work has shown that Cel6B has an erratic processive run length on the cellulose surface, with the majority of its time being paused, making the enzyme prone to denaturation when stuck on the surface during processive catalytic turnover cycles.^27^ With the higher binding affinity towards cellulose seen for positively charged CBMs, it is plausible the D5 CBM2a construct may be more prone to getting stuck, and with salt present to screen some electrostatics, those pauses may be shortened causing the slight higher activity with the addition of salt. Still, it is likely that this improvement is more structure-function related with D5 CBM2a – WT Cel6B resisting denaturation when stuck on the surface. It is also interesting to note the improvements are clear on cellulose only and not biomass, and thus, the choice of pretreatment technology is equally as important for implementation when building a synergistic cocktail. An alternative pretreatment like dilute acid pretreatment may be more favorable as it will produce a biomass substrate more similar to cellulose-I used here, thus providing optimal substrate for similarly engineered enzymes.

### Thermodynamic decoupling between domains boosts intrinsic thermostability for best performer

Based on the observed differences in the Cel6B library in pH and salt-dependent assays, we next assessed a potential shift in thermostability with protein thermal shift assays. Previous work with supercharged endocellulase Cel5A showed no significant improvement in melting temperatures. In almost every case tested as well, two individual peaks were identified in melt curves signifying individual melting between the CBM and CD for supercharged constructs while the native endocellulase showed coupled melting with CBM and CD.^37^ Here, melt curves were constructed using differential scanning fluorimetry (DSF) for the CBM2a – Cel6B chimeras to identify if any structural stabilization resulting from supercharging impacted the thermostability of these enzymes (**Figure 4**).

**Figure 4.**
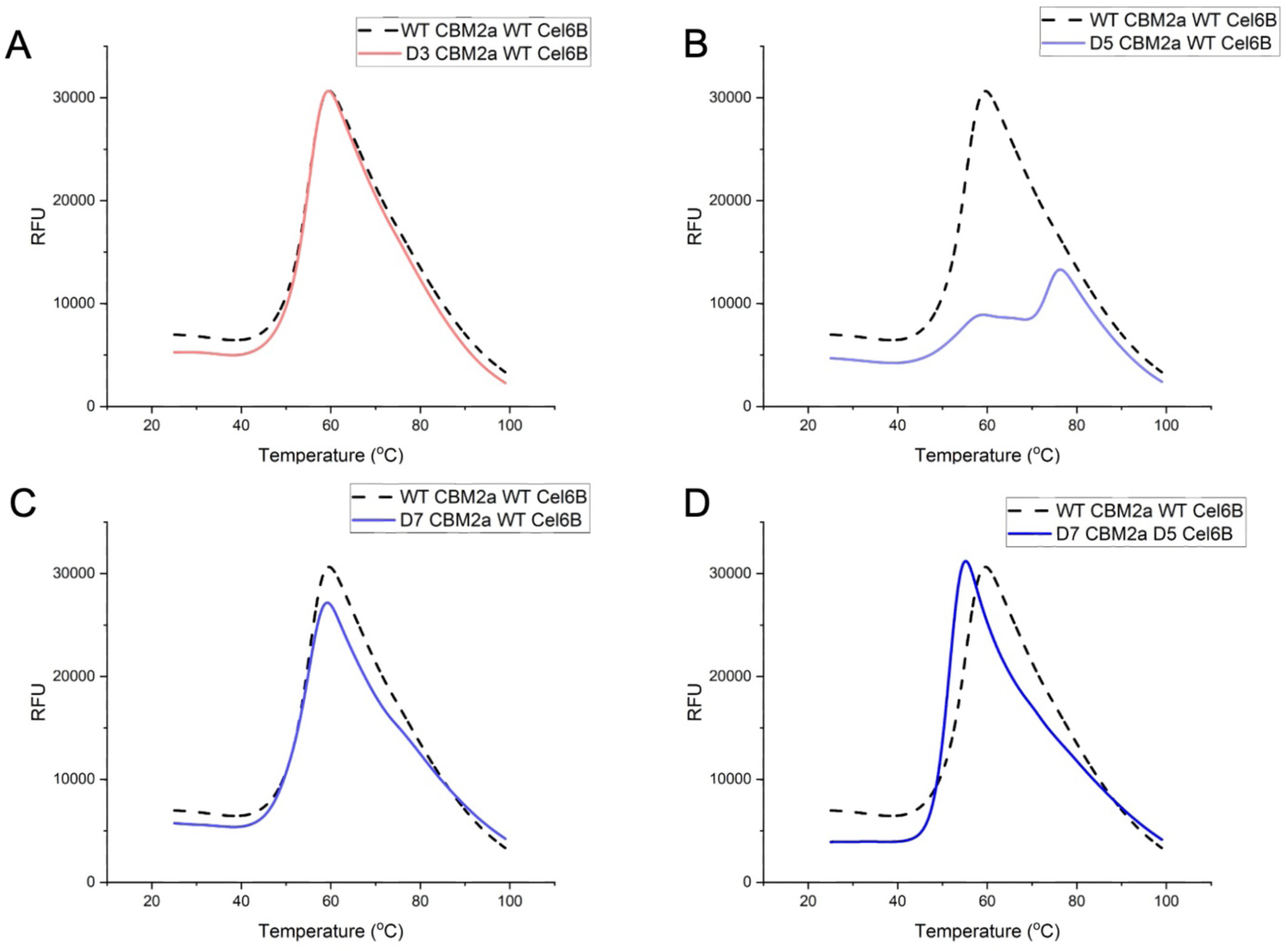
Thermodynamic decoupling of CBM and catalytic-domain melting in D5 CBM2a–WT Cel6B retains catalytic-domain thermostability. Enzyme melt curves comparing the native enzyme (dashed black line) to purified (A) D3 CBM2a – WT Cel6B, (B) D5 CBM2a – WT Cel6B, (C) D7 CBM2a – WT Cel6B, and (D) D7 CBM2a – D5 Cel6B. Thermal shift assays were conducted using 5 µL of 50x SYPRO orange dye, 2.5 µL of 1M sodium phosphate pH 6.0, 17.5 µL of DI water, and 25 µL of 10 µM protein dilution (in DI water) in an optically clear 96-well PCR plate. Heat denaturation and subsequent fluorescent signal monitoring were done in a Bio-Rad CFX Opus 96 real-time PCR system by equilibrating samples first to 25 °C and heating to 99 °C with a 0.5 °C increment and 5 second read time. All proteins were tested in quadruplicates and data was processed in MATLAB to average replicates, subtract background fluorescence of a water blank, and smooth curves.

For all constructs except D5 CBM2a – WT Cel6B (**Figure 4B**), a single cooperative unfolding transition was observed corresponding to the coupled melting of the smaller CBM initiating melting of the catalytic domain as well. This pattern is essentially unchanged in the negatively charged D3 CBM2a–Cel6B (**Figure 4A**) and the highly positive D7 CBM2a–Cel6B (**Figure 4C**) mutants. The positively charged combinatorial construct D7 CBM2a – D5 Cel6B (**Figure 4D**) shows a slight leftward shift indicating destabilization and a lower melt temperature, but still only one melt transition is seen clearly. D5 CBM2a – WT Cel6B is the peculiar case where the unfolding of the D5 CBM does not trigger cooperative unfolding of Cel6B. Although the CBM melts at roughly the same temperature as in the other constructs, the catalytic domain of D5 CBM2a–WT Cel6B remains folded until about 70 °C, a rightward shift of roughly 18 °C compared to the native enzyme, where cooperative melting occurs at much lower temperatures. Melting temperatures were estimated by taking the negative derivative of the melt curves and are depicted in **Figure 5**. The D3 CBM2a – WT Cel6B and D7 CBM2a – WT Cel6B chimeras show no significant change in reported melt temperature compared to the native enzyme (**Figure 5 A, C**) whereas the combinatorial construct D7 CBM2a – D5 Cel6B (**Figure 5D**) showed a near 4-degree decrease in melting temperature. For the best performer, D5 CBM2a – WT Cel6B (**Figure 5B**), the first melt transition occurs around the same temperature as all other constructs, with the second melt temperature around 73.45 °C, more akin to the expected melting temperature of enzymes from thermophiles.

**Figure 5.**
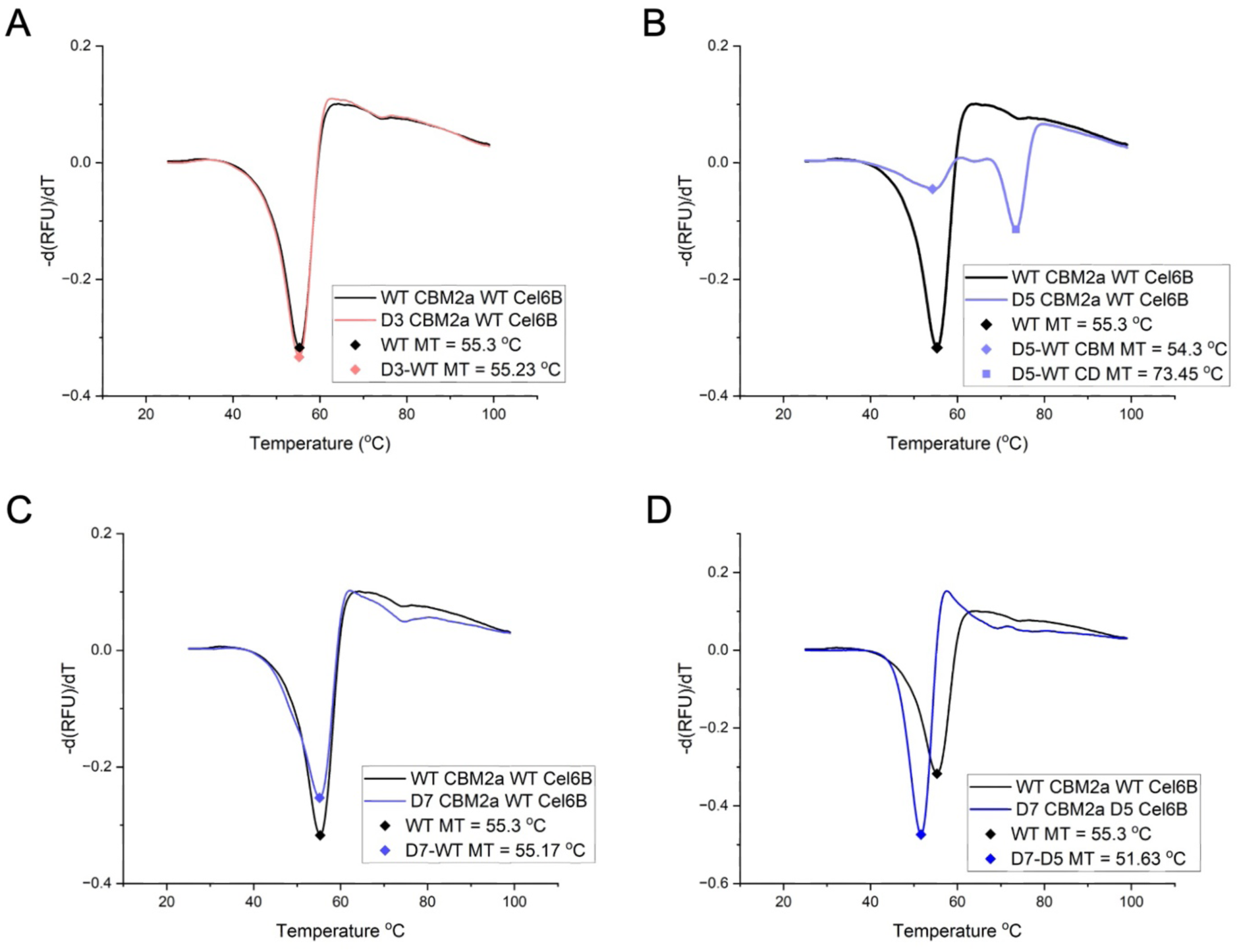
Derivative melt curves reveal melting temperature for native and supercharged Cel6B enzymes. Derivative melt plots comparing the native CBM2a – Cel6B (black dashed line) to (A) D3 CBM2a – WT Cel6B, (B) D5 CBM2a – WT Cel6B, (C) D7 CBM2a – WT Cel6B, and (D) D7 CBM2a – D5 Cel6B. Melting temperatures were estimated by taking the negative derivative of melt curve data and finding the minimum value of the resulting plot. Data was processed in MATLAB by first averaging the four replicates per melt curve and subtracting background fluorescence measured with a water blank. The negative derivative was calculated using the MATLAB gradient function and minima were identified with the min function in MATLAB.

This distinction underscores that enhanced performance can arise not only from stabilizing specific domains but also from tuning how domains interact and unfold under thermal stress. This coupling between domains was observed with the native endocellulase CBM2a – Cel5A, but the CBM and Cel5A CD melted together at the intrinsic melting temperature of the catalytic domain. For many supercharged constructs, especially those positively supercharged, including one chimera with only a positive supercharged CBM and native CD, this decoupling between domain melting points was observed.^37^ For Cel5A, this decoupling became a limitation because the full-length enzyme was originally melting around 60-70 °C. For Cel6B, the clear limitation is that the coupled melting between domains is initiated at the CBM melting point and not higher CD melting point like for Cel5A. Due to this, the enzyme denatures completely at the lower melting point, thus, decoupling becomes advantageous because, although CBM function is lost, the CD remains intact and able to continue catalysis. This is especially critical for an exocellulase like Cel6B that is frequently dwelled on the surface of cellulose where unfolding will occur.^27^ Therefore, the improvements observed for D5 CBM2a – WT Cel6B as a result of supercharging likely do not stem from improved substrate binding like with Cel5A, but instead improved mechanistic function as a result from this thermodynamic decoupling.

### Supercharged cellulases that perform best individually are the best synergistic partners

Having identified D5 CBM2a–WT Cel6B as a superior exocellulase mutant for cellulose hydrolysis and established the driving force behind this improvement, the next question was whether this enzyme would synergize with supercharged endocellulase variants. To test this, D5 CBM2a – WT Cel6B was paired with several top-performing Cel5A variants and compared to the native enzyme. First, individual enzyme activity was assessed on AFEX corn stover and cellulose-I for each supercharged construct and both native full-length Cel5A and Cel6B enzymes at a constant enzyme loading (**Figure 6**). The most notable observation is the clear substrate preference between the exocellulases and endocellulases. The Cel5A constructs exhibit much higher activity on biomass than Cel6B, likely due to the multifunctional nature of this endocellulase, which is also active on xylan.^37^ However, both the native Cel6B and D5 CBM construct show higher activity on crystalline cellulose, which is to be expected for an exocellulase. The positively supercharged D3 CBM2a–Cel5A endocellulase outperforms the other Cel5A variants on cellulose I, likely due to its optimized electrostatic complementarity with the cellulose surface, though this advantage hinders performance when lignin is present.

**Figure 6.**
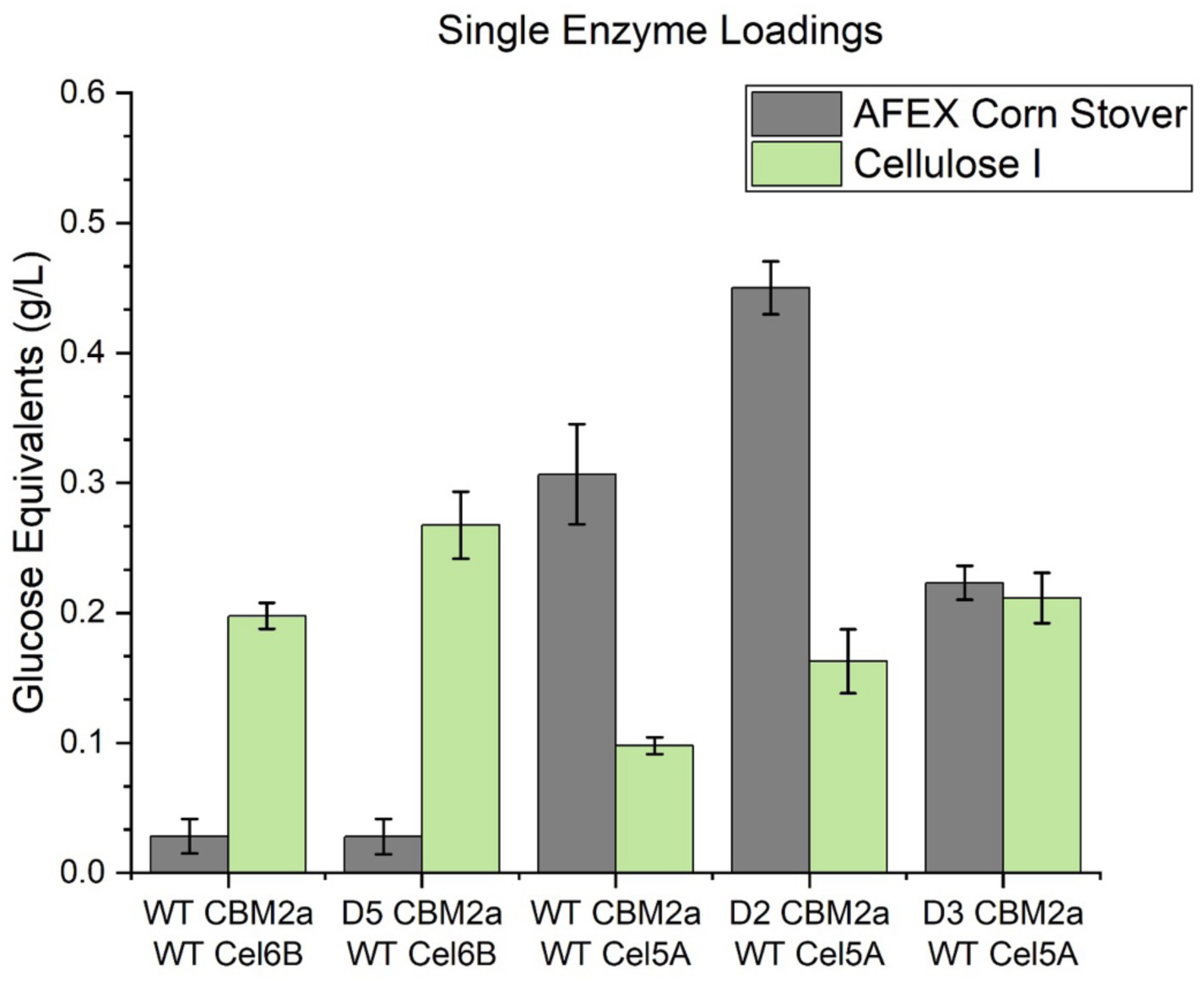
Activity of native exocellulase Cel6B and endocellulase Cel5A compared to best performing supercharged variants. The activity of the listed constructs was tested on a constant 2 mg of either AFEX pretreated corn stover or crystalline cellulose-I with a constant loading of 120 nmol enzyme per gram substrate. Both substrates were prepared as slurries in DI water to a concentration of 25 g/L for AFEX and 100 g/L for cellulose-I. Activity was tested in 50 mM sodium phosphate pH 6.0 based on the optimal hydrolysis pH identified from previous assays and all dilutions were made in DI water. Assay plates were incubated for 24 hours at 60 °C with 5 RPM end-over-end mixing and reducing sugar equivalents was estimated using DNS assay and compared to glucose standards. A buffer blank was included and to subtract background absorbance resulting from potential soluble reducing sugars present in the substrate that may potentially give a response in the DNS assay. Data points represent the average of four replicates and error bars represent the standard deviation from the mean.

Each exo:endo enzyme pair was tested across several ratios, with a 1:3 endo:exo ratio expected to be optimal based on prior work with *T. fusca* cellulases (**Figure 7**).^23^ With AFEX corn stover however, there is little improvement when adding Cel6B (native or D5 CBM construct) compared to the lone endocellulase treatment. This is especially true for the D2 CBM2a – Cel5A endocellulase that showed high individual activity on AFEX corn stover. This further emphasizes the importance of pretreatment chemistry when selecting enzymes for biorefinery applications. AFEX preserves the overall crystallinity of cellulose and retains a higher concentration of amorphous cellulose and hemicellulose^11,12,56^ producing a much more endocellulase friendly substrate. In contrast, a pretreatment method like DA pretreatment restructures the crystalline network in cellulose, decreasing the overall dispersion of amorphous regions and removing hemicellulose, thus retaining a much more exo friendly crystalline substrate.^14,57^ Exo-endo synergism is highly substrate dependent, with both crystallinity and bulk substrate properties impacting the overall effectiveness of exo-endo pairs.^58^

**Figure 7.**
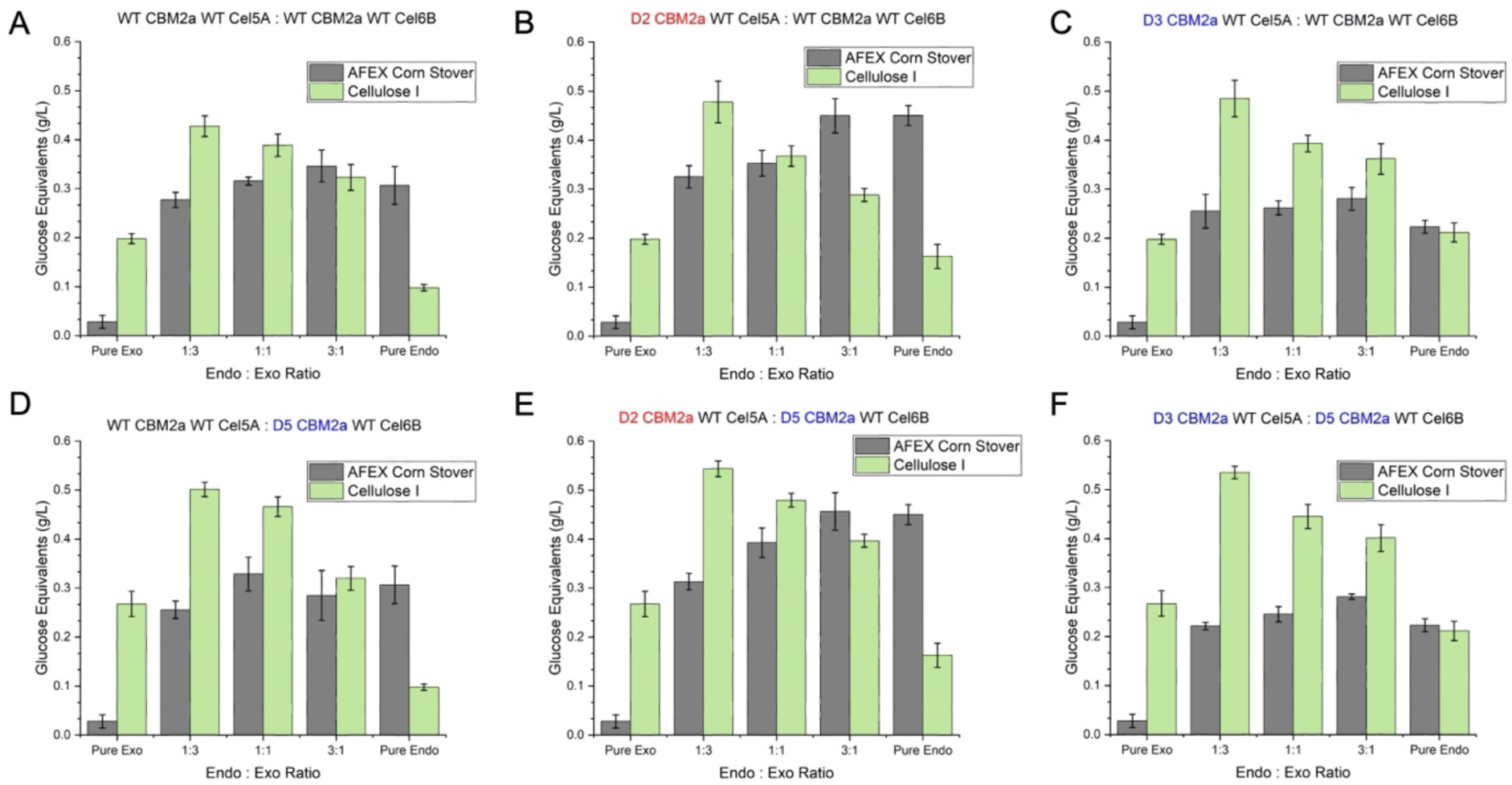
Supercharged cellulases with the highest individual catalytic efficiencies form the most synergistic enzyme pairs. Endo-exo synergistic hydrolysis of AFEX corn stover and cellulose I by the WT CBM2a – WT Cel6B exocellulase with (A) WT CBM2a – WT Cel5A (B) D2 CBM2a – WT Cel5A, and (C) D3 CBM2a – WT Cel5A endocellulase, and the D5 CBM2a – WT Cel6B exocellulase with (D) WT CBM2a – WT Cel5A, (E) D2 CBM2a – WT Cel5A, and (F) D3 CBM2a – WT Cel5A endocellulase. Enzyme activity was tested on AFEX corn stover and crystalline cellulose – I with a total amount of 2 mg of substrate prepared as a 25 g/L (AFEX) or 100 g/L (cellulose – I) slurry in DI water. A total enzyme loading of 120 nmol enzyme per gram of substrate was used, and to keep this loading constant, different endo:exo ratios adjusted the load of cellulase so the total amount of enzyme loaded was 120 nmol/g at each different ratio. Hydrolysis was conducted in 50 mM sodium phosphate buffer pH 6.0 for 24 hours at 60 °C with 5 RPM end-over-end mixing. Reducing sugar equivalents were estimated via DNS assay and compared to glucose standards. Data points represent the average of four replicates and error bars represent standard deviation from the mean.

A 1:3 endo:exo ratio did show the highest overall hydrolysis yield on crystalline cellulose – I for each pair tested, and in every case, a higher overall yield than each individual construct tested alone indicating synergistic hydrolysis with the enzyme pair. The highest hydrolysis yield on crystalline cellulose is observed when pairing D5 CBM2a – Cel6B with D2 CBM2a – WT Cel5A (**Figure 7E**). Interestingly, pairing oppositely charged enzymes did not produce any inhibitory interactions with oppositely charged domains that negatively impacted hydrolysis yield. A similar, yet slightly lower, activity was observed when pairing D5 CBM2a – Cel6B with D3 CBM2a – WT Cel5A, two positively charged enzyme (**Figure 7F**). For this positive – positive pairing though, a low AFEX corn stover activity was observed, stemming from the poorer activity of the positive endocellulase with lignin present. Overall, synergy scaled with the individual performance of each enzyme. The results suggest that synergism does not arise from charge complementarity or increased colocalization, but from the intrinsic catalytic efficiency of the individual enzymes. Instead, it appears that individual enzyme performance is the strongest predictor of overall synergistic gain through cellulase supercharging as those showing the highest isolated activities produce the best pairing.

## CONCLUSION

In this study we have successfully improved the activity of a GH family-6 processive exocellulase and its fused native family-2a carbohydrate binding module from the thermophile *Thermobifida fusca* using protein supercharging as computational design strategy. A total of 16 supercharged constructs were computationally identified and supercharged, eight per domain, and a library of 32 fusion constructs was generated and systematically screened. Activity was first assessed from soluble cell lysates to identify a key construct: D5 CBM2a – WT Cel6B that exhibits a higher hydrolytic activity on all substrates tested near the published pH optimum of 5.5. This chimera showed near a 2.3-fold increase in hydrolysis yield on crystalline cellulose compared to the native enzyme when purified and exhibit an interesting shift in its melt curve that helps reveal the intrinsic melting temperature of the native thermophilic catalytic domain. When pairing with a supercharged endocellulase from previous studies, both highest performing supercharged endo- and exo-cellulases, respectively, proved to be the best pairing for synergistic cellulose degradation. The performance of this supercharged exocellulase showed surprising differences compared to the endocellulase Cel5A. For an endocellulase, supercharging heavily influenced adsorption and desorption to and from the cellulose surface. The much simpler endocellulase mechanism seems to be more limited in these steps where imparting positive electrostatic charges complimentary to the negative substrate surface assists in adsorption to an extent, with the Sabatier principle being a major design consideration to ensure intermediate strength binding interactions are optimized. This was not the case for Cel6B. It seems that the exocellulase mechanism is less hindered by initial binding to the substrate. Although removing a CBM from Cel6B will drastically drop overall activity,^27,59,60^ it appears there is limited benefit in further tuning surface characteristics for Cel6B in particular. What is more nuanced with this enzyme is the dwell time it spends on the cellulose surface which seems to be more limiting in the overall function of Cel6B. Cel6B spends majority of its processive cycle paused on the cellulose surface where either desorption or denaturation will occur. This is where the D5 CBM2a – WT Cel6B mutant seems to clearly shine, as thermodynamic decoupling of the melting points of the CBM and CD allow it to retain its folded catalytic core even at higher temperatures over extended durations of catalysis. A similar trend was observed for Cel5A where the CBM and CD melted together but decoupled in supercharged constructs. For Cel5A, this wasn’t an issue since the tandem melting occurred at the higher melting temperature of the Cel5A CD. For native Cel6B and majority of its supercharged constructs, denaturation occurs at the lower melting temperature of the CBM for the full-length enzyme, but by decoupling domain melting, this issue is mitigated for D5 CBM2a – WT Cel6B.

Ultimately, for implementation, a full synergistic cocktail of enzymes is required for effective utilization of pretreated biomass deconstruction to fermentable sugars. The work here provides insight into two key points: (i) the importance of curtailing substrate pretreatment to make biomass most amenable to enzymatic deconstruction, and (ii) how cellulase pairs can be produced for improved synergistic deconstruction. Regarding the first point, the real improvements of D5 CBM2a – WT Cel6B were observed on crystalline cellulose – I while activity on AFEX showed similar results compared to the native enzyme. Implementation of this enzyme would likely require alternative substrate pretreatment, such as dilute acid pretreatment, where a higher proportion of crystalline cellulose remains compared to amorphous regions and hemicellulose for better utilization. Regarding enzyme pairs, we have shown here that supercharging is a successful method for improving not only the activity of an endocellulase like Cel5A, but an exocellulase like Cel6B as well. This is critical for the implementation of this technology because current biorefineries are hindered by low enzyme activity and high loading requirements that drive up processing costs.^19^ The 2.3-fold improvements in hydrolysis yields observed here allows for a meaningful impact in the economic feasibility of implementation. Technoeconomic analyses have shown that improvements in enzyme activity to the scale observed here can result in $0.25 – 0.30 per gallon decrease in minimum ethanol selling price.^15^

The exo-endo pair used here that have shown to be an improved synergistic pair of cellulases for lignocellulosic deconstruction originate from the same thermophilic microbe *T. fusca*. Engineering and improving enzymes from the same microbe is a positive step towards designing a complete synergistic cocktail of *T. fusca* enzymes for biorefinery implementation for various pretreatment technologies. This microbe contains a full set of CAZymes required for lignocellulose valorization,^23^ and improving a minimum set of enzymes from this microbe can facilitate integrated enzyme production in a future biorefinery scheme further improving the profitability of such a process.^61^ Still, more work needs to be done to improve the activity of other celluloytic and hemicelluloytic enzymes from *T. fusca* which is a logical next step towards implementation. However, the work describe here shows much promise in generating enzymes for improved biomass utilization with supercharging. As we have observed, supercharging is an effective tool for improving enzyme activity that can generate better synergistic partners. The results affirm the guiding principle of this study: *there is an “I” in team*—individual improvements in designed enzymes directly translate to higher cooperative performance seen during biomass hydrolysis.

## Supporting information

SI Sequences and Data

## ACKNOWLEDGEMENTS

This study was primarily supported by NSF CBET awards (1604421 and 1846797) and Sungrant award (#S005193-USDA). ADC was supported by the Biotechnology Training Program fellowship that was funded by Rutgers University, Sungrant award, and the National Institute of General Medical Sciences of the National Institutes of Health under award number T32 GM135141. All genes used for expression of recombinant constructs were provided by the Department of Energy Joint Genome Institute (DOE-JGI) supported by the Community Science Program Gene Synthesis Award (CSP-503631 Syn). We would like to thank Deanne Sammond (NREL) for help with computational design of supercharged cellulases at the onset of the DOE-JGI award. Finally, we would like to also thank Professor Rebecca Garlock-Ong (MTU) and Professor Bruce Dale (MSU) for kindly providing access to AFEX treated corn stover used in this study.

## Notes

### Competing Interest Statement

The authors have declared no competing interest.

## REFERENCES

(1) Chundawat, S. P. S.; Beckham, G. T.; Himmel, M.; Dale, B. E. Deconstruction of Lignocellulosic Biomass to Fuels and Chemicals. Annu. Rev. Chem. Biomol. Eng. 2011, 2, 121–145. 10.1146/annurev-chembioeng-061010-114205.

(2) Tuck, C. O.; Pérez, E.; Horváth, I. T.; Sheldon, R. A.; Poliakoff, M. Valorization of Biomass: Deriving More Value from Waste. Science. 2012, pp 695–699. 10.1126/science.1218930.

(3) Somerville, C.; Youngs, H.; Taylor, C.; Davis, S. C.; Long, S. P. Feedstocks for Lignocellulosic Biofuels. Science. 2010. 10.1126/science.1189268.

(4) Payne, C. M.; Knott, B. C.; Mayes, H. B.; Hansson, H.; Himmel, M. E.; Sandgren, M.; Ståhlberg, J.; Beckham, G. T. Fungal Cellulases. Chemical Reviews. 2015. 10.1021/cr500351c.

(5) Somerville, C.; Bauer, S.; Brininstool, G.; Facette, M.; Hamann, T.; Milne, J.; Osborne, E.; Paredez, A.; Persson, S.; Raab, T.; Vorwerk, S.; Youngs, H. Toward a Systems Approach to Understanding Plant Cell Walls. Science. 2004. 10.1126/science.1102765.

(6) Himmel, M. E.; Ding, S. Y.; Johnson, D. K.; Adney, W. S.; Nimlos, M. R.; Brady, J. W.; Foust, T. D. Biomass Recalcitrance: Engineering Plants and Enzymes for Biofuels Production. Science. 2007. 10.1126/science.1137016.

(7) Jeoh, T.; Ishizawa, C. I.; Davis, M. F.; Himmel, M. E.; Adney, W. S.; Johnson, D. K. Cellulase Digestibility of Pretreated Biomass Is Limited by Cellulose Accessibility. Biotechnol. Bioeng. 2007, 98 (1), 112–122. 10.1002/bit.21408.

(8) Zeng, Y.; Zhao, S.; Yang, S.; Ding, S. Y. Lignin Plays a Negative Role in the Biochemical Process for Producing Lignocellulosic Biofuels. Current Opinion in Biotechnology. 2014. 10.1016/j.copbio.2013.09.008.

(9) McCann, M. C.; Carpita, N. C. Biomass Recalcitrance: A Multi-Scale, Multi-Factor, and Conversion-Specific Property. Journal of Experimental Botany. 2015. 10.1093/jxb/erv267.

(10) da Costa Sousa, L.; Chundawat, S. P.; Balan, V.; Dale, B. E. “Cradle-to-Grave” Assessment of Existing Lignocellulose Pretreatment Technologies. Current Opinion in Biotechnology. 2009. 10.1016/j.copbio.2009.05.003.

(11) Balan, V.; Bals, B.; Chundawat, S. P. S.; Marshall, D.; Dale, B. E. Lignocellulosic Biomass Pretreatment Using AFEX. Methods in molecular biology (Clifton, N.J.). 2009. 10.1007/978-1-60761-214-8_5.

(12) Chundawat, S. P. S.; Pal, R. K.; Zhao, C.; Campbell, T.; Teymouri, F.; Videto, J.; Nielson, C.; Wieferich, B.; Sousa, L.; Dale, B. E.; Balan, V.; Chipkar, S.; Aguado, J.; Burke, E.; Ong, R. G. Ammonia Fiber Expansion (AFEX) Pretreatment of Lignocellulosic Biomass. J. Vis. Exp. 2020, 2020 (158). 10.3791/57488.

(13) Da Costa Sousa, L.; Jin, M.; Chundawat, S. P. S.; Bokade, V.; Tang, X.; Azarpira, A.; Lu, F.; Avci, U.; Humpula, J.; Uppugundla, N.; Gunawan, C.; Pattathil, S.; Cheh, A. M.; Kothari, N.; Kumar, R.; Ralph, J.; Hahn, M. G.; Wyman, C. E.; Singh, S.; Simmons, B. A.; Dale, B. E.; Balan, V. Next-Generation Ammonia Pretreatment Enhances Cellulosic Biofuel Production. Energy Environ. Sci. 2016, 9 (4). 10.1039/c5ee03051j.

(14) Shekiro, J.; Kuhn, E. M.; Nagle, N. J.; Tucker, M. P.; Elander, R. T.; Schell, D. J. Characterization of Pilot-Scale Dilute Acid Pretreatment Performance Using Deacetylated Corn Stover. Biotechnol. Biofuels 2014, 7 (1). 10.1186/1754-6834-7-23.

(15) Humbird, D.; Davis, R.; Tao, L.; Kinchin, C.; Hsu, D.; Aden, A.; Schoen, P.; Lukas, J.; Olthof, B.; Worley, M.; Sexton, D.; Dudgeon, D. Process Design and Economics for Biochemical Conversion of Lignocellulosic Biomass to Ethanol: Dilute-Acid Pretreatment and Enzymatic Hydrolysis of Corn Stover. In NREL/TP-5100-47764; National Renewable Energy Laboratory, Golden, CO, USA, 2011.

(16) Gao, D.; Haarmeyer, C.; Balan, V.; Whitehead, T. A.; Dale, B. E.; Chundawat, S. P. Lignin Triggers Irreversible Cellulase Loss during Pretreated Lignocellulosic Biomass Saccharification. Biotechnol. Biofuels 2014, 7 (1), 175. 10.1186/s13068-014-0175-x.

(17) Davis, R.; Bartling, A. *Biochemical Conversion of Lignocellulosic Biomass to Hydrocarbon Fuels and Products: 2022 State of Technology and Future Research*. Golden, CO: National Renewable Energy Laboratory. Technical Report NREL/TP-5100-85469; 2023.

(18) Scown, C. D.; Baral, N. R.; Yang, M.; Vora, N.; Huntington, T. Technoeconomic Analysis for Biofuels and Bioproducts. Curr. Opin. Biotechnol. 2021, 67, 58–64. 10.1016/j.copbio.2021.01.002.

(19) Klein-Marcuschamer, D.; Oleskowicz-Popiel, P.; Simmons, B. A.; Blanch, H. W. The Challenge of Enzyme Cost in the Production of Lignocellulosic Biofuels. Biotechnol. Bioeng. 2012, 109 (4), 1083–1087. 10.1002/bit.24370.

(20) Luo, X.; Liu, J.; Zheng, P.; Li, M.; Zhou, Y.; Huang, L.; Chen, L.; Shuai, L. Promoting Enzymatic Hydrolysis of Lignocellulosic Biomass by Inexpensive Soy Protein. Biotechnol. Biofuels 2019, 12 (1), 1–13. 10.1186/s13068-019-1387-x.

(21) Yang, B.; Wyman, C. E. BSA Treatment to Enhance Enzymatic Hydrolysis of Cellulose in Lignin Containing Substrates. Biotechnol. Bioeng. 2006, 94 (4), 611–617. 10.1002/bit.20750.

(22) Nemmaru, B.; DeChellis, A.; Patil, N.; Chundawat, S. P. S. CAZyme Characterization and Engineering for Biofuels Applications. In Handbook of Biorefinery Research and Technology; 2023. 10.1007/978-94-007-6724-9_32-1.

(23) Wilson, D. B. Studies of Thermobifida Fusca Plant Cell Wall Degrading Enzymes. Chem. Rec. 2004, 4 (2). 10.1002/tcr.20002.

(24) Sandgren, M.; Wu, M.; Karkehabadi, S.; Mitchinson, C.; Kelemen, B. R.; Larenas, E. A.; Ståhlberg, J.; Hansson, H. The Structure of a Bacterial Cellobiohydrolase: The Catalytic Core of the Thermobifida Fusca Family GH6 Cellobiohydrolase Cel6B. J. Mol. Biol. 2013, 425 (3), 622–635. 10.1016/j.jmb.2012.11.039.

(25) Chundawat, S. P. S.; Nemmaru, B.; Hackl, M.; Brady, S. K.; Hilton, M. A.; Johnson, M. M.; Chang, S.; Lang, M. J.; Huh, H.; Lee, S. H.; Yarbrough, J. M.; López, C. A.; Gnanakaran, S. Molecular Origins of Reduced Activity and Binding Commitment of Processive Cellulases and Associated Carbohydrate-Binding Proteins to Cellulose III. J. Biol. Chem. 2021, 296. 10.1016/j.jbc.2021.100431.

(26) Wu, M.; Bu, L.; Vuong, T. V.; Wilson, D. B.; Crowley, M. F.; Sandgren, M.; Ståhlberg, J.; Beckham, G. T.; Hansson, H. Loop Motions Important to Product Expulsion in the Thermobifida Fusca Glycoside Hydrolase Family 6 Cellobiohydrolase from Structural and Computational Studies. J. Biol. Chem. 2013, 288 (46), 33107–33117. 10.1074/jbc.M113.502765.

(27) Johnson, M. M.; DeChellis, A.; Nemmaru, B.; Chundawat, S. P. S.; Lang, M. J. Thermobifida Fusca Cel6B Moves Bidirectionally While Processively Degrading Cellulose. Biotechnol. biofuels Bioprod. 2024, 17 (1), 140. 10.1186/s13068-024-02588-0.

(28) Alasepp, K.; Borch, K.; Cruys-Bagger, N.; Badino, S.; Jensen, K.; Sørensen, T. H.; Windahl, M. S.; Westh, P. In Situ Stability of Substrate-Associated Cellulases Studied by DSC. Langmuir 2014, 30 (24). 10.1021/la500161e.

(29) Heinzelman, P.; Snow, C. D.; Smith, M. A.; Yu, X.; Kannan, A.; Boulware, K.; Villalobos, A.; Govindarajan, S.; Minshull, J.; Arnold, F. H. SCHEMA Recombination of a Fungal Cellulase Uncovers a Single Mutation That Contributes Markedly to Stability. J. Biol. Chem. 2009, 284 (39). 10.1074/jbc.C109.034058.

(30) Heinzelman, P.; Komor, R.; Kanaan, A.; Romero, P.; Yu, X.; Mohler, S.; Snow, C.; Arnold, F. Efficient Screening of Fungal Cellobiohydrolase Class i Enzymes for Thermostabilizing Sequence Blocks by SCHEMA Structure-Guided Recombination. Protein Eng. Des. Sel. 2010, 23 (11). 10.1093/protein/gzq063.

(31) Heinzelman, P.; Snow, C. D.; Wu, I.; Nguyen, C.; Villalobos, A.; Govindarajan, S.; Minshull, J.; Arnold, F. H.; Designed, F. H. A.; Performed, C. N. A Family of Thermostable Fungal Cellulases Created by Structure-Guided Recombination; 2009; Vol. 106.

(32) Wu, I.; Arnold, F. H. Engineered Thermostable Fungal Cel6A and Cel7A Cellobiohydrolases Hydrolyze Cellulose Efficiently at Elevated Temperatures. Biotechnol. Bioeng. 2013, 110 (7). 10.1002/bit.24864.

(33) Vuong, T. V.; Wilson, D. B. Processivity, Synergism, and Substrate Specificity of Thermobifida Fusca Cel6B. Appl. Environ. Microbiol. 2009, 75 (21). 10.1128/AEM.01260-09.

(34) Lawrence, M. S.; Phillips, K. J.; Liu, D. R. Supercharging Proteins Can Impart Unusual Resilience. J. Am. Chem. Soc. 2007, 129 (33). 10.1021/ja071641y.

(35) Haarmeyer, C. N.; Smith, M. D.; Chundawat, S. P. S.; Sammond, D.; Whitehead, T. A. Insights into Cellulase-Lignin Non-Specific Binding Revealed by Computational Redesign of the Surface of Green Fluorescent Protein. Biotechnol. Bioeng. 2017, 114 (4). 10.1002/bit.26201.

(36) Whitehead, T. A.; Bandi, C. K.; Berger, M.; Park, J.; Chundawat, S. P. S. Negatively Supercharging Cellulases Render Them Lignin-Resistant. ACS Sustain. Chem. Eng. 2017, 5 (7), 6247–6252. 10.1021/acssuschemeng.7b01202.

(37) DeChellis, A.; Nemmaru, B.; Sammond, D.; Douglass, J.; Patil, N.; Reste, O.; Chundawat, S. P. S. Supercharging Carbohydrate Binding Module Alone Enhances Endocellulase Thermostability, Binding, and Activity on Cellulosic Biomass. ACS Sustain. Chem. Eng. 2024, 12 (9). 10.1021/acssuschemeng.3c06266.

(38) Zhang, X.; Qu, T.; Mosier, N. S.; Han, L.; Xiao, W. Cellulose Modification by Recyclable Swelling Solvents. Biotechnol. Biofuels 2018, 11 (1). 10.1186/s13068-018-1191-z.

(39) Song, Y.; Dimaio, F.; Wang, R. Y. R.; Kim, D.; Miles, C.; Brunette, T.; Thompson, J.; Baker, D. High-Resolution Comparative Modeling with RosettaCM. Structure 2013, 21 (10), 1735–1742.

(40) Lawrence, M. S.; Phillips, K. J.; Liu, D. R. Supercharging Proteins Can Impart Unusual Resilience. J. Am. Chem. Soc. 2007, 129 (33), 10110–10112. 10.1021/ja071641y.

(41) Der, B. S.; Kluwe, C.; Miklos, A. E.; Jacak, R.; Lyskov, S.; Gray, J. J.; Georgiou, G.; Ellington, A. D.; Kuhlman, B. Alternative Computational Protocols for Supercharging Protein Surfaces for Reversible Unfolding and Retention of Stability. PLoS One 2013, 8 (5). 10.1371/journal.pone.0064363.

(42) Studier, F. W. Protein Production by Auto-Induction in High Density Shaking Cultures. Protein Expr. Purif. 2005, 41 (1), 207–234. 10.1016/J.PEP.2005.01.016.

(43) Liu, Y.; Nemmaru, B.; Chundawat, S. P. S. Thermobif Ida f Usca Cellulases Exhibit Increased Endo−Exo Synergistic Activity, but Lower Exocellulase Activity, on Cellulose-III. 2020. 10.1021/acssuschemeng.9b06792.

(44) Miller, G. L. Use of Dinitrosalicylic Acid Reagent for Determination of Reducing Sugar. Anal. Chem. 1959, 31 (3). 10.1021/ac60147a030.

(45) Mayer, M. P.; Bukau, B. Hsp70 Chaperones: Cellular Functions and Molecular Mechanism. Cellular and Molecular Life Sciences. 2005. 10.1007/s00018-004-4464-6.

(46) Bu, L.; Beckham, G. T.; Shirts, M. R.; Nimlos, M. R.; Adney, W. S.; Himmel, M. E.; Crowley, M. F. Probing Carbohydrate Product Expulsion from a Processive Cellulase with Multiple Absolute Binding Free Energy Methods. J. Biol. Chem. 2011, 286 (20), 18161–18169. 10.1074/jbc.M110.212076.

(47) Murphy, L.; Bohlin, C.; Baumann, M. J.; Olsen, S. N.; Sørensen, T. H.; Anderson, L.; Borch, K.; Westh, P. Product Inhibition of Five Hypocrea Jecorina Cellulases. Enzyme Microb. Technol. 2013, 52 (3). 10.1016/j.enzmictec.2013.01.002.

(48) Olsen, J. P.; Alasepp, K.; Kari, J.; Cruys-Bagger, N.; Borch, K.; Westh, P. Mechanism of Product Inhibition for Cellobiohydrolase Cel7A during Hydrolysis of Insoluble Cellulose. Biotechnol. Bioeng. 2016, 113 (6). 10.1002/bit.25900.

(49) Wilks, J. C.; Slonczewski, J. L. PH of the Cytoplasm and Periplasm of Escherichia Coli: Rapid Measurement by Green Fluorescent Protein Fluorimetry. J. Bacteriol. 2007, 189 (15). 10.1128/JB.00615-07.

(50) Le Costaouëc, T.; Pakarinen, A.; Várnai, A.; Puranen, T.; Viikari, L. The Role of Carbohydrate Binding Module (CBM) at High Substrate Consistency: Comparison of Trichoderma Reesei and Thermoascus Aurantiacus Cel7A (CBHI) and Cel5A (EGII). Bioresour. Technol. 2013, 143. 10.1016/j.biortech.2013.05.079.

(51) Várnai, A.; Siika-Aho, M.; Viikari, L. Carbohydrate-Binding Modules (CBMs) Revisited: Reduced Amount of Water Counterbalances the Need for CBMs. Biotechnol. Biofuels 2013, 6 (1). 10.1186/1754-6834-6-30.

(52) Brady, S. K.; Sreelatha, S.; Feng, Y.; Chundawat, S. P. S.; Lang, M. J. Cellobiohydrolase 1 from Trichoderma Reesei Degrades Cellulose in Single Cellobiose Steps. Nat. Commun. 2015, 6 (May), 10149. 10.1038/ncomms10149.

(53) Sabatier, P. Hydrogénations et Déshydrogénations Par Catalyse. Berichte der Dtsch. Chem. Gesellschaft 1911, 44 (3), 1984–2001. 10.1002/CBER.19110440303.

(54) Kari, J.; Olsen, J. P.; Jensen, K.; Badino, S. F.; Krogh, K. B. R. M.; Borch, K.; Westh, P. Sabatier Principle for Interfacial (Heterogeneous) Enzyme Catalysis. ACS Catal. 2018, 8 (12). 10.1021/acscatal.8b03547.

(55) Arnling Bååth, J.; Jensen, K.; Borch, K.; Westh, P.; Kari, J. Sabatier Principle for Rationalizing Enzymatic Hydrolysis of a Synthetic Polyester. JACS Au 2022, 2 (5). 10.1021/jacsau.2c00204.

(56) Chundawat, S. P. S.; Donohoe, B. S.; Da, L.; Sousa, C.; Elder, T.; Agarwal, U. P.; Lu, F.; Ralph, J.; Himmel, M. E.; Balan, V.; Dale, B. E. Multi-Scale Visualization and Characterization of Lignocellulosic Plant Cell Wall Deconstruction during Thermochemical Pretreatment † ‡. 10.1039/c0ee00574f.

(57) Pu, Y.; Hu, F.; Huang, F.; Davison, B. H.; Ragauskas, A. J. Assessing the Molecular Structure Basis for Biomass Recalcitrance during Dilute Acid and Hydrothermal Pretreatments. Biotechnol. Biofuels 2013, 6 (1). 10.1186/1754-6834-6-15.

(58) Kostylev, M.; Wilson, D. Synergistic Interactions in Cellulose Hydrolysis. Biofuels. 2012. 10.4155/bfs.11.150.

(59) Herve, C.; Rogowski, A.; Blake, A. W.; Marcus, S. E.; Gilbert, H. J.; Knox, J. P. Carbohydrate-Binding Modules Promote the Enzymatic Deconstruction of Intact Plant Cell Walls by Targeting and Proximity Effects. Proc. Natl. Acad. Sci. 2010, 107 (34), 15293–15298. 10.1073/pnas.1005732107.

(60) Reyes-Ortiz, V.; Heins, R. A.; Cheng, G.; Kim, E. Y.; Vernon, B. C.; Elandt, R. B.; Adams, P. D.; Sale, K. L.; Hadi, M. Z.; Simmons, B. A.; Kent, M. S.; Tullman-Ercek, D. Addition of a Carbohydrate-Binding Module Enhances Cellulase Penetration into Cellulose Substrates. Biotechnol. Biofuels 2013, 6 (1). 10.1186/1754-6834-6-93.

(61) Johnson, E. Integrated Enzyme Production Lowers the Cost of Cellulosic Ethanol. *Biofuels*, Bioprod. Biorefining 2016, 10 (2). 10.1002/bbb.1634.

