## Supplementary material for "There is an “I” in team: individual improvements in supercharged cellulase cocktail facilitates cooperative cellulose degradation": SI Figures and Tables

**Number of SI pages in current PDF (SI-Supplementary Information)**: 24

**Number of SI figures in current PDF (SI-Supplementary Information):** 5

**Number of SI tables in current PDF (SI-Supplementary Information):** 18

**Number of SI files (including current PDF):** 2

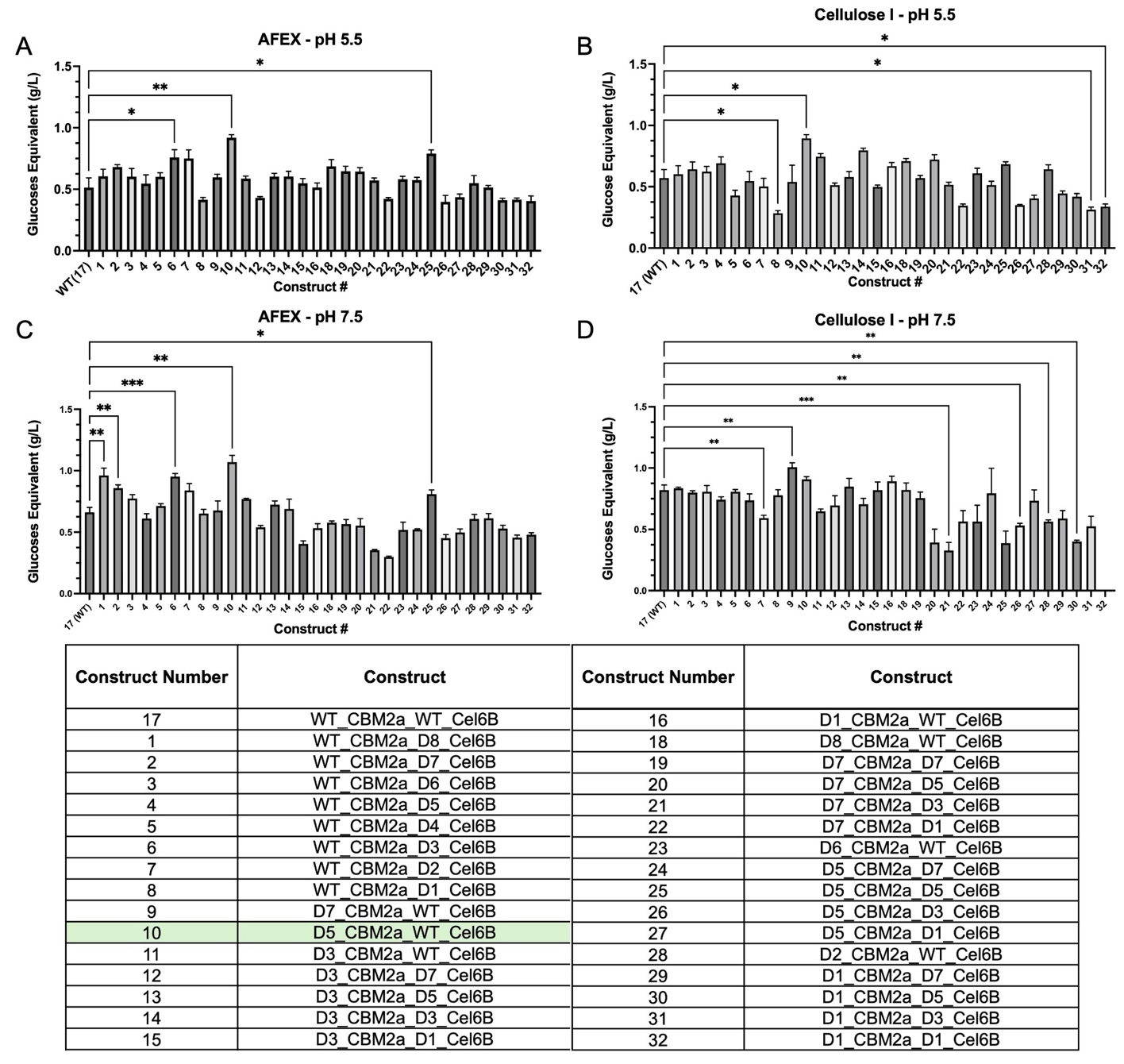
**Supplementary Figure S1. Statistical analysis of cell lysate screen on insoluble cellulosic substrates.** Data shown here is the same as depicted in Figure 2 of the main manuscript. The numbering scheme for constructs in S1 reflects the original numbering used in culturing, but this was reorganized for clarity in the final manuscript. As was depicted in the main manuscript, data represents the average of four technical replicates and error bars depict standard deviation from the mean. PASC data was omitted from the present figure because several groups showed statistically significant differences and thus only the tabulated statistics were included here and are viewable in the subsequent tables. Statistical significance between each construct was analyzed using Welch’s ANOVA for normally distributed data with unequal variances (determined from both Levene’s test and Bartlett’s test) in Prism 10 software and yielded a significant difference for each data set (p < 0.05). Differences in the means for each construct was compared to the native enzyme using Dunnett’s post hoc analysis and these tables are available in this SI appendix. Statistical significance as per Dunnett’s analysis is indicated in **Figure S1** by the following: * for p ≤ 0.05, ** for p ≤ 0.01, *** for p ≤ 0.001, and **** for p ≤ 0.0001.

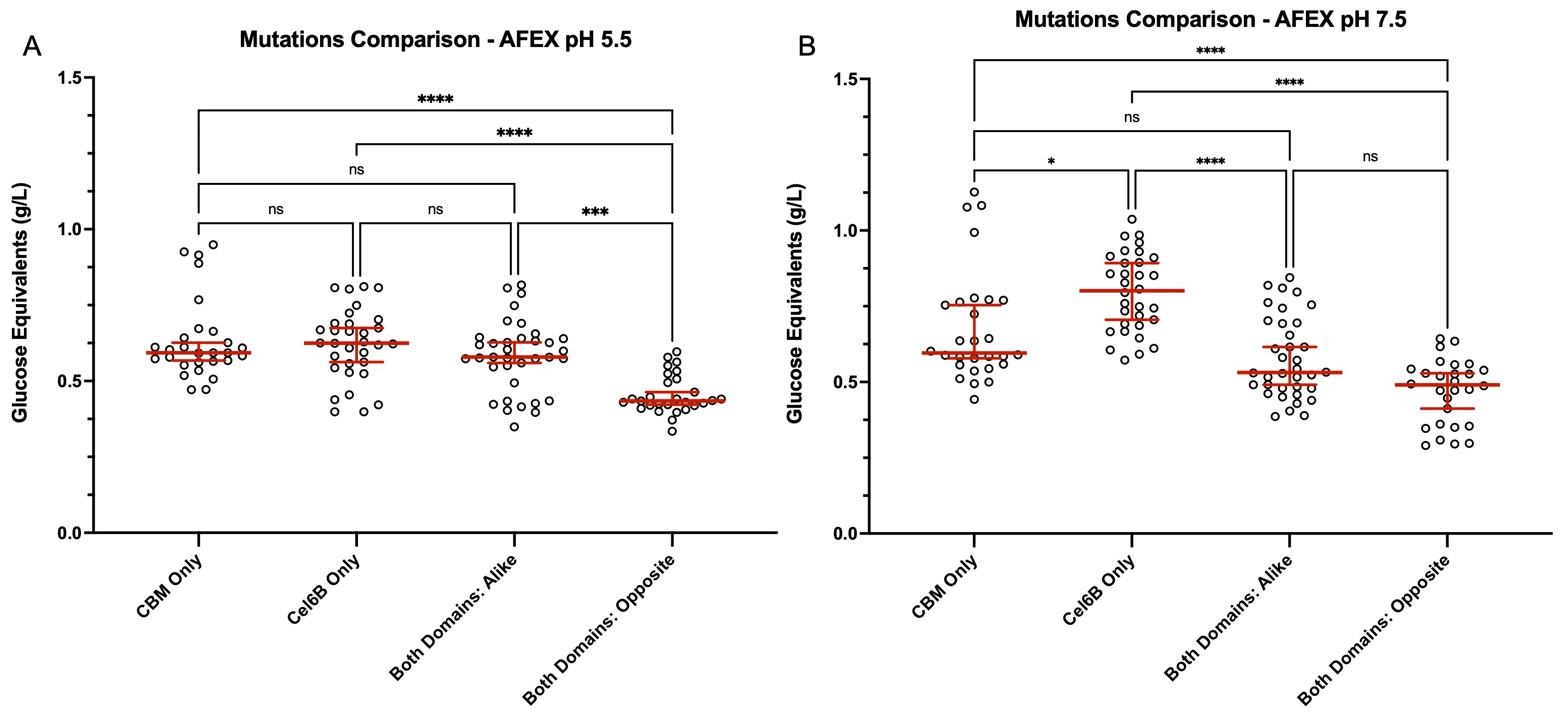
**Supplementary Figure S2. Comparing mutation placement with activity on AFEX corn stover.** Data depicted utilizes lysate screen activity on AFEX corn stover at (A) pH 5.5 and (B) pH 7.5. Bins correlate to full-length CBM2a – Cel6B constructs that contain only a supercharged CBM, only a supercharged Cel6B, two supercharged domains that are alike (positive – positive or negative – negative), and two supercharged domains that are opposite in charge (negative – positive or positive – negative). Grouping of the data set in this manner produces non-parametric data based on both a Shapiro-Wilk’s and D’Agostino test. A Kruskal – Wallis ANOVA was used to compare statistically significant differences from the median (non-parametric) and Dunn’s multiple comparison was used for post hoc analysis. Individual points correspond to one technical replicate with horizontal red lines showing the median of the entire group and error bars depict 95% confidence intervals. All statistical analysis and plotting were done in Prism 10 software. Statistical significance is indicated by the following notation: ns for no significant difference, * for p ≤ 0.05, ** for p ≤ 0.01, *** for p ≤ 0.001, and **** for p ≤ 0.0001.

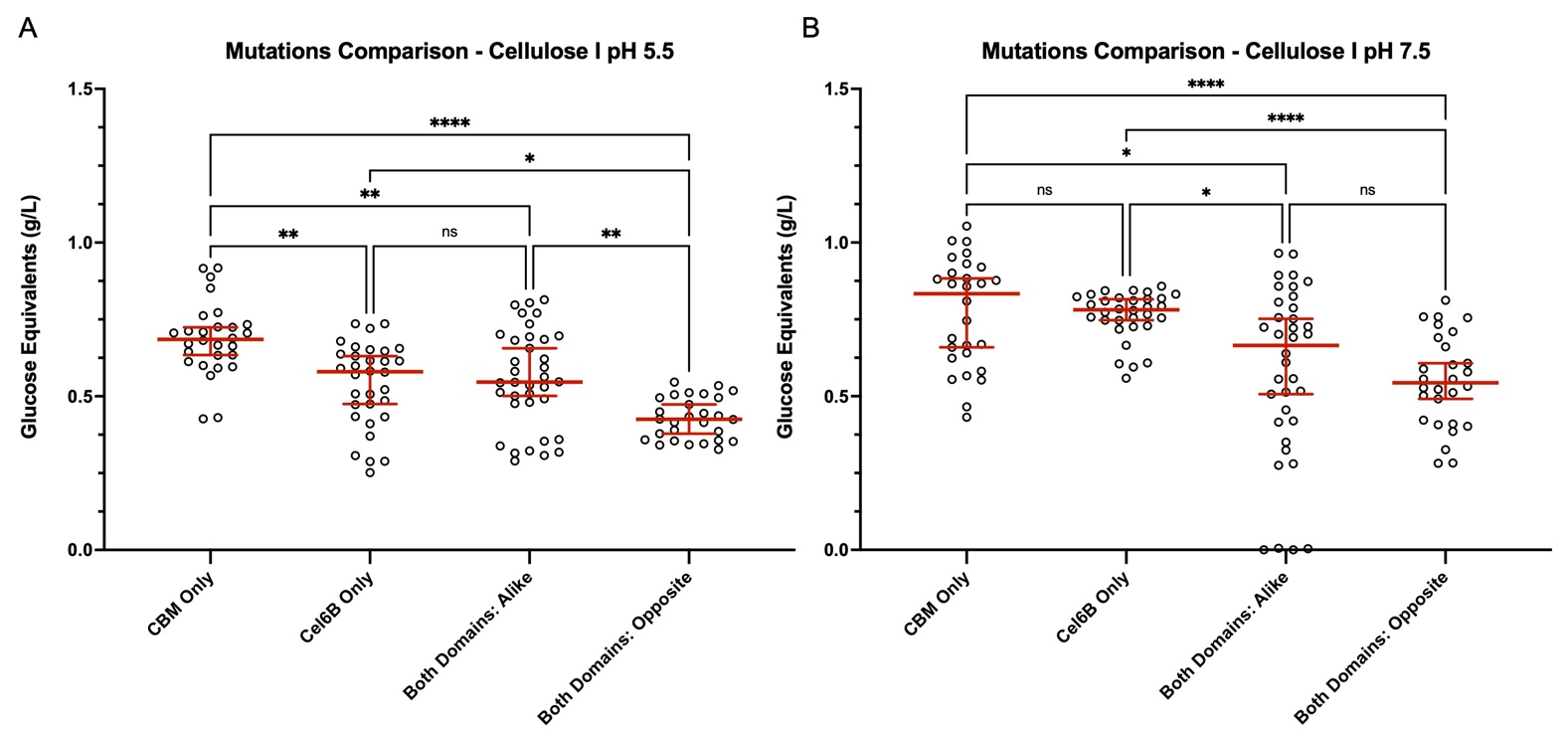

**Supplementary Figure S3. Comparing mutation placement with activity on crystalline cellulose – I.** Data depicted utilizes lysate screen activity on cellulose – I at (A) pH 5.5 and (B) pH 7.5. Bins correlate to full-length CBM2a – Cel6B constructs that contain only a supercharged CBM, only a supercharged Cel6B, two supercharged domains that are alike (positive – positive or negative – negative), and two supercharged domains that are opposite in charge (negative – positive or positive – negative). Grouping of the data set in this manner produces non-parametric data based on both a Shapiro-Wilk’s and D’Agostino test. A Kruskal – Wallis ANOVA was used to compare statistically significant differences from the median (non-parametric) and Dunn’s multiple comparison was used for post hoc analysis. Individual points correspond to one technical replicate with horizontal red lines showing the median of the entire group and error bars depict 95% confidence intervals. All statistical analysis and plotting were done in Prism 10 software. Statistical significance is indicated by the following notation: ns for no significant difference, * for p ≤ 0.05, ** for p ≤ 0.01, *** for p ≤ 0.001, and **** for p ≤ 0.0001.

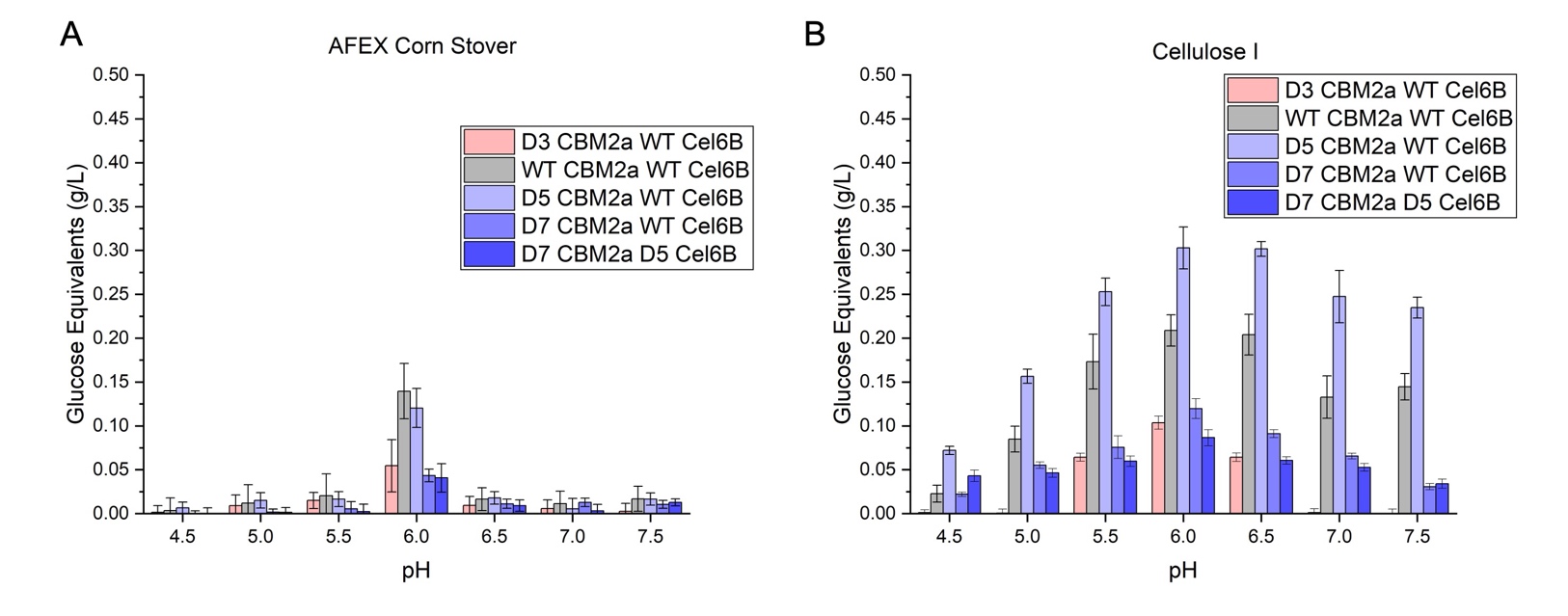
**Supplementary Figure S3. Purified enzyme pH sweep shows no shift in optimal hydrolysis pH.** Hydrolysis of (A) AFEX corn stover and (B) crystalline cellulose – I for several purified CBM2a – Cel6B constructs tested at several different solution pH. A constant substrate amount of 2 mg was used in all reactions and substrate stocks were prepared as a 25 g/L slurry for AFEX and 100 g/L for cellulose – I in DI water. Enzyme activity was assessed at a constant 120 nmol of enzyme per gram of substrate, and all reactions were conducted in 0.2 mL round bottom microplates. Solution pH was controlled by adding 10 µL of 1 M stock sodium acetate (pH 4.5 – 5.5) or sodium phosphate (pH 6.0 – 7.5) buffer for an effective buffer concentration of 50 mM. Plates were sealed with a TPE capmat-96 microplate seal and covered with clear packing tape to avoid any evaporative losses. Plates were incubated at 60 °C for 24 hours with 5 RPM end-over-end mixing in a VWR hybridization oven. Reducing sugar equivalents were quantified by analyzing the soluble hydrolysate post incubation by DNS assay and compared to glucose standards. All data points correspond to the average of four replicates and error bars denote standard deviation from the mean. Constructs are color coded based on net charge, ranging from negative (red) to most positive (dark blue). The native enzyme is represented in gray.

**Supplementary Figure S5. A clear Sabatier optimum likely does not exist for supercharged exocellulase Cel6B.**
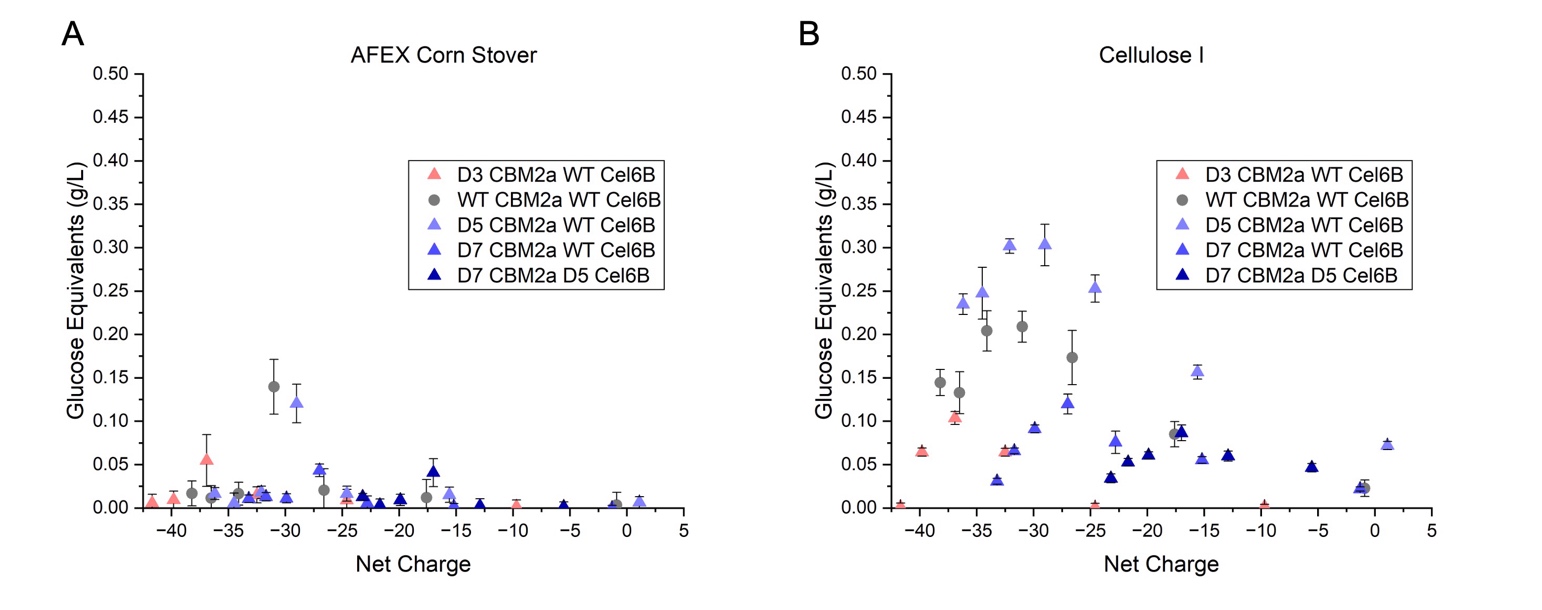
 Correlating activity at different pH on (A) AFEX corn stover and (B) cellulose – I to enzyme net charge. Data depicted in figure S4 was used and correlated to the full-length enzyme net charge at each pH tested as calculated by Prot pi online tool (protpi.ch/Calculator/ProteinTool) based on the primary amino acid sequence for each construct. His-tags were not included in the sequences when calculating charge since they are assumed to be a constant contributor to each enzyme, and thus, only differences based on supercharging were considered. Since no shift in enzyme optimal pH was seen for the exocellulase, there is no clear correlation to enzyme charge and activity like was seen with a supercharged endocellulase (Cel5A) in our previous study. Each data point represents the average of four technical replicates and error bars denote standard deviation from the mean. Constructs are color coded based on net charge, ranging from negative (red) to most positive (dark blue). The native enzyme is represented in gray.

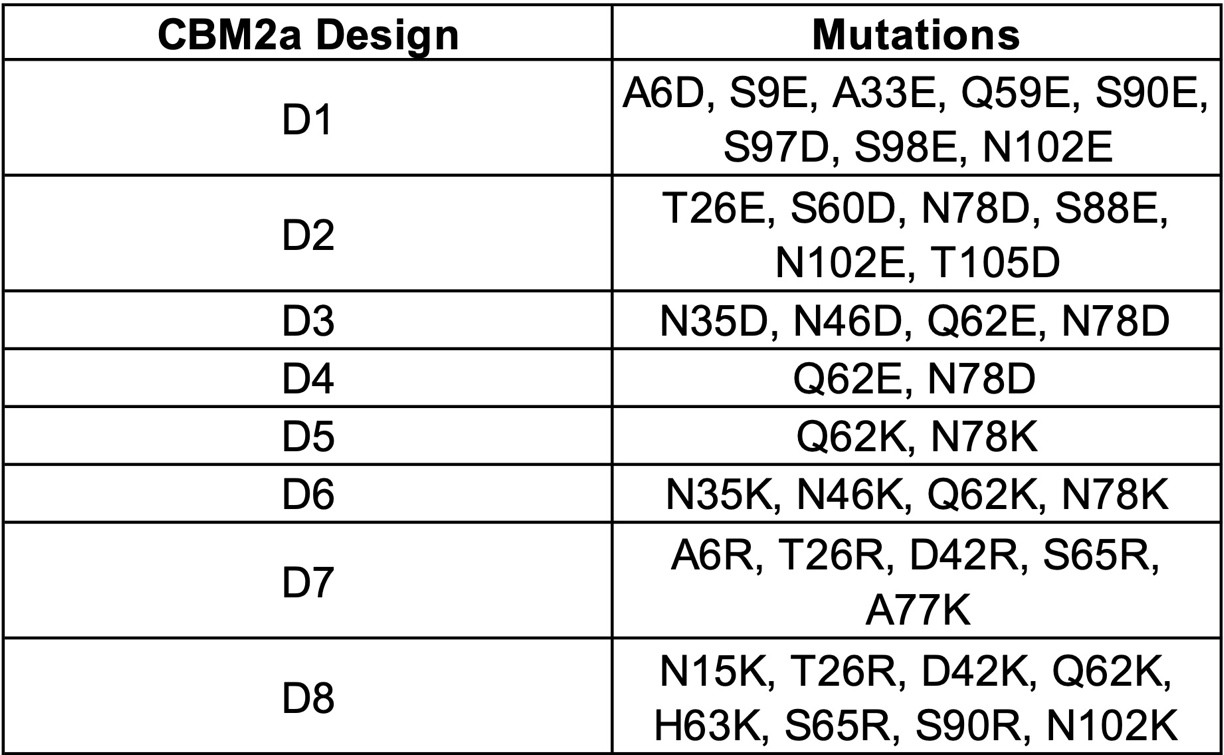

**Supplementary Table T1. Mutations made on wildtype CBM2a to generate supercharged mutants.** Tabulated list of each mutation made to the native CBM to produce the CBM2a design denoted in column 1. Constructs D1 – D4 are negatively supercharged and D5 – D8 are positively supercharged.

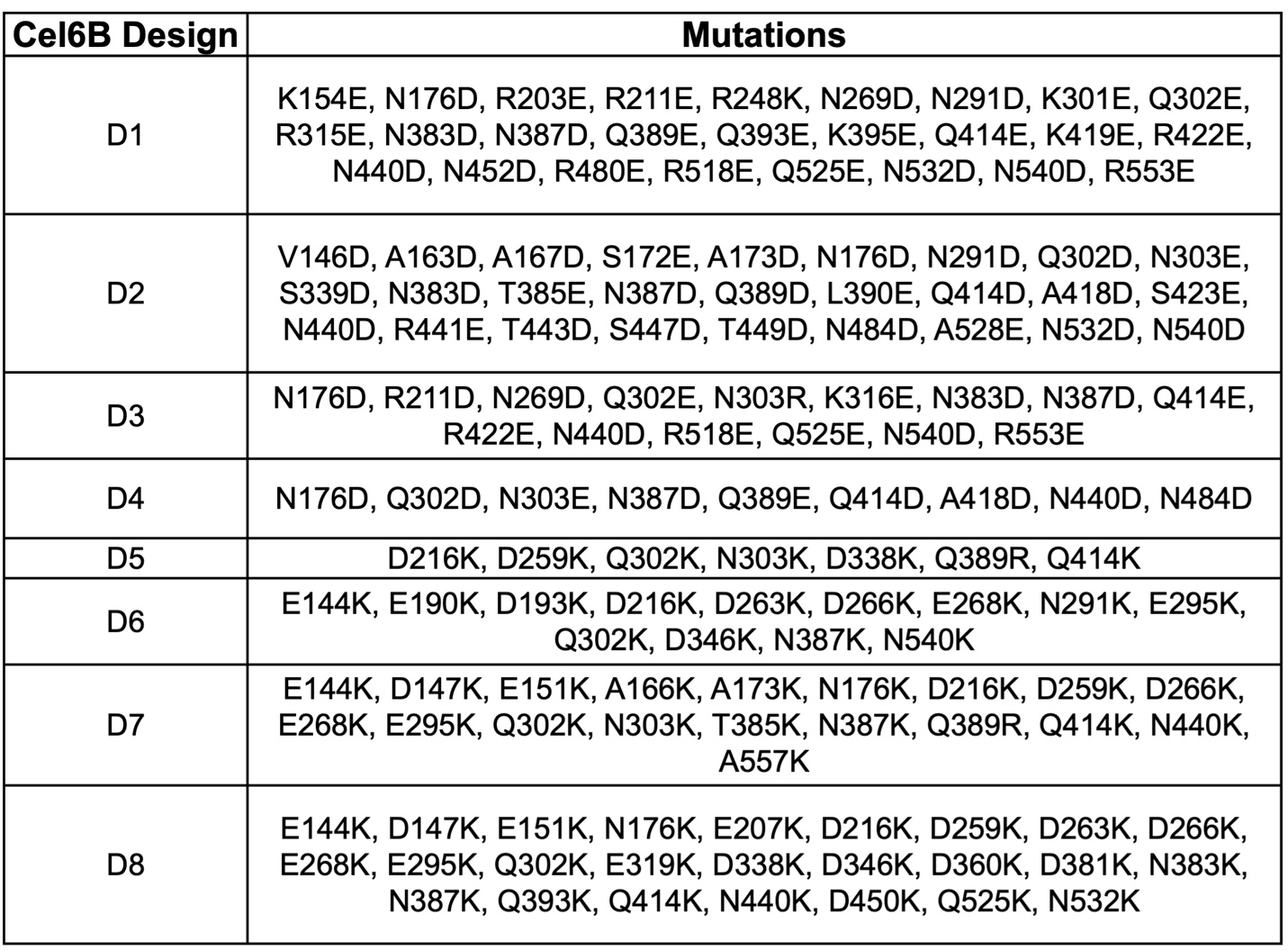
**Supplementary Table T2. Mutations on wildtype Cel6B to generate supercharged mutants.** Tabulated list of each mutation made to the native Cel6B CD to produce the CD design denoted in column 1. Constructs D1 – D4 are negatively supercharged and D5 – D8 are positively supercharged.

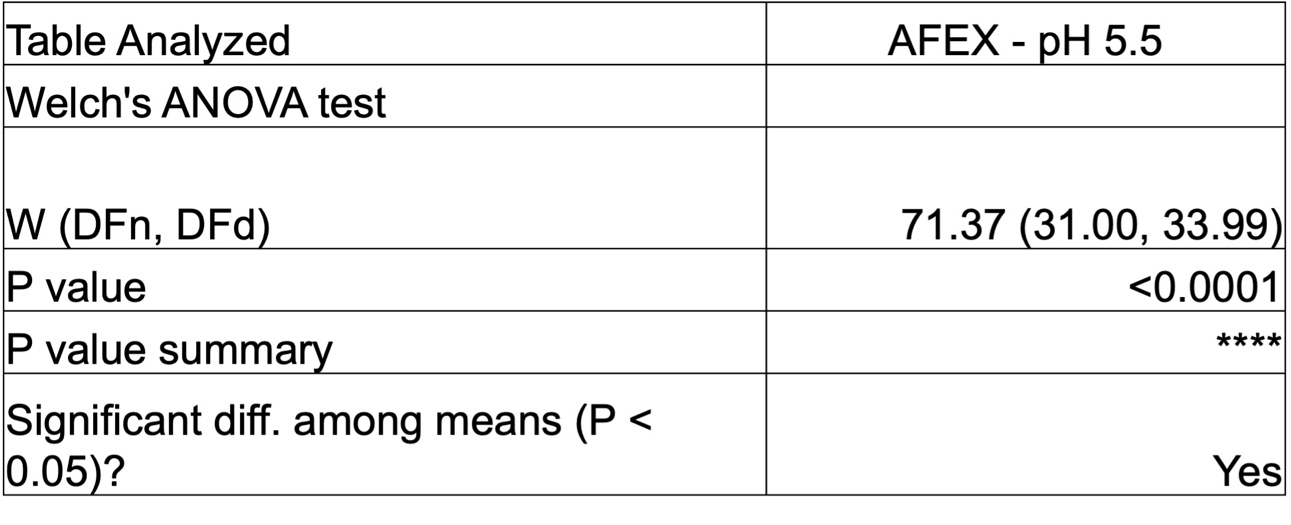

**Supplementary Table T3. Welch’s ANOVA result for AFEX corn stover lysate screen data at pH 5.5.** Anova result depicted indicates a statistically significant difference (p ≤ 0.05) in mean activity for the supercharged library. This statistical analysis corresponds to the data shown in **Figure 2A** of the main manuscript and **Figure S1A** of the SI appendix.

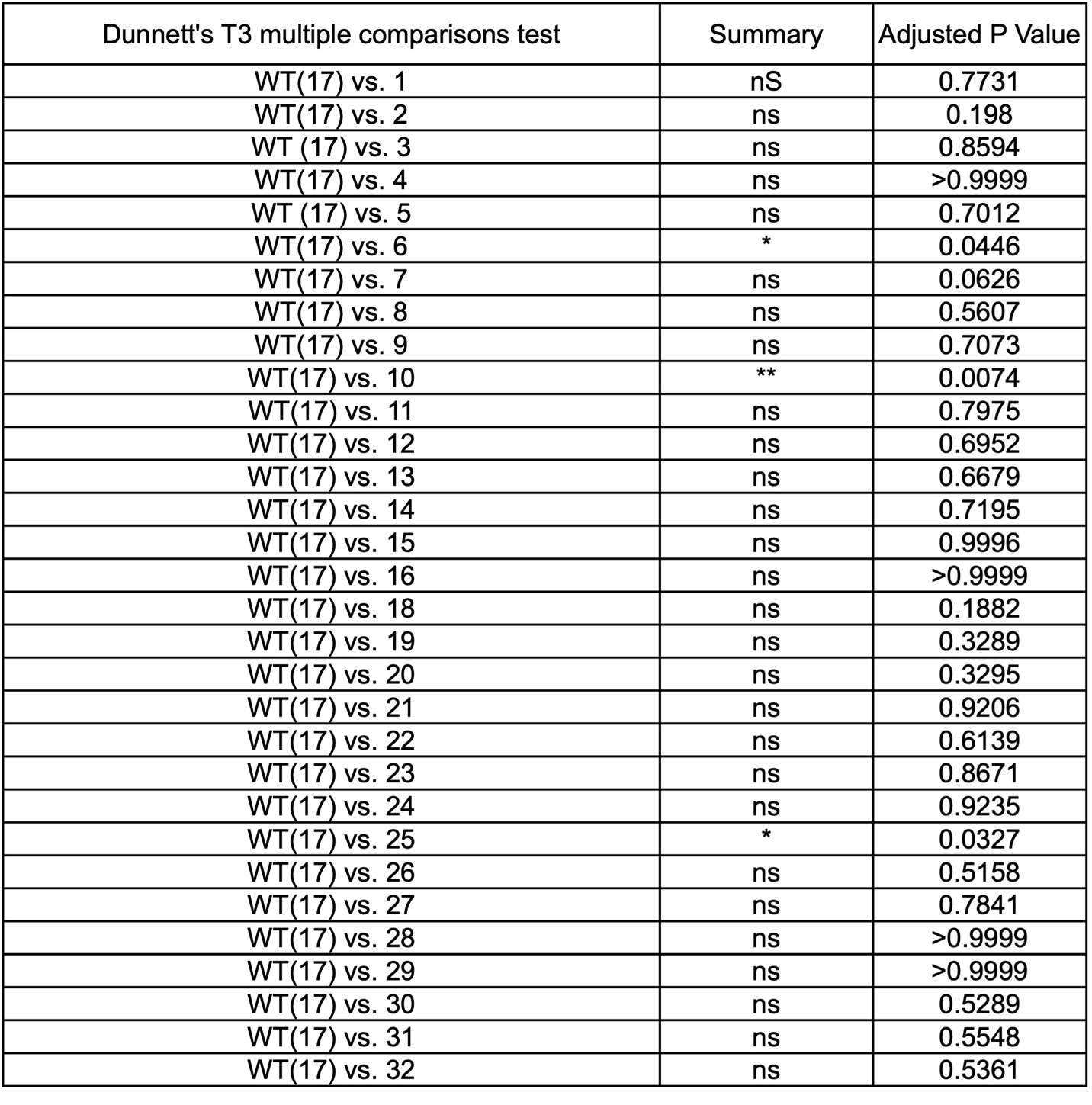

**Supplementary Table T4. Dunnett’s post hoc analysis for AFEX corn stover lysate screen data at pH 5.5.** Post hoc follow-up analysis after Welch’s ANOVA comparing the mean activity for each supercharged construct to the native enzyme. This statistical analysis corresponds to the data shown in **Figure 2A** of the main manuscript and **Figure S1A** of the SI appendix.

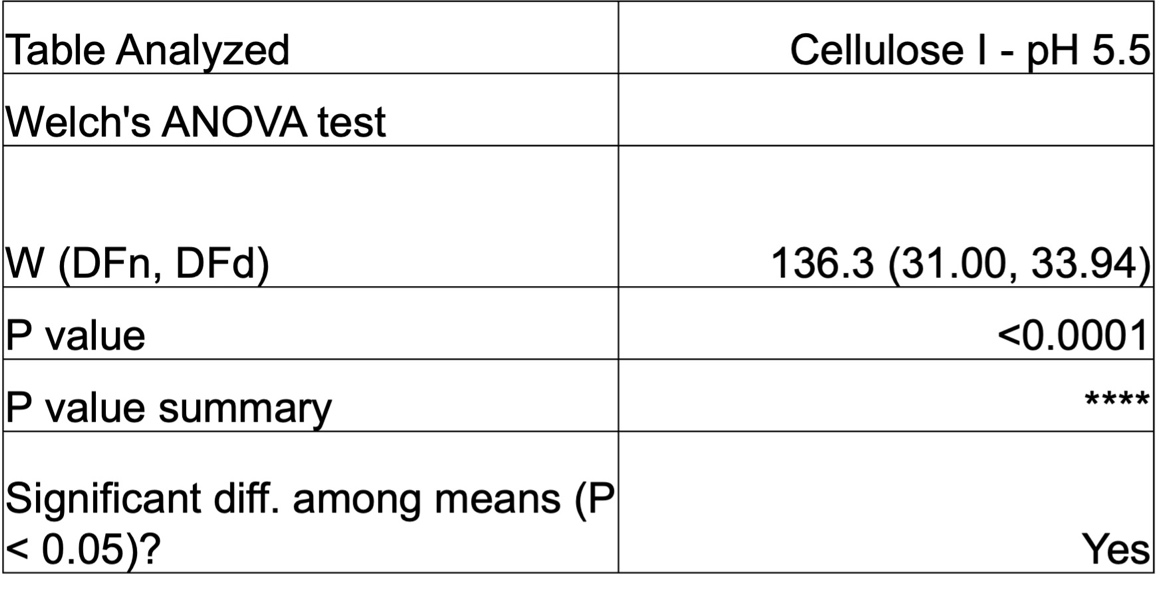

**Supplementary Table T5. Welch’s ANOVA result for cellulose – I lysate screen data at pH 5.5.** Anova result depicted indicates a statistically significant difference (p ≤ 0.05) in mean activity for the supercharged library. This statistical analysis corresponds to the data shown in **Figure 2B** of the main manuscript and **Figure S1B** of the SI appendix.

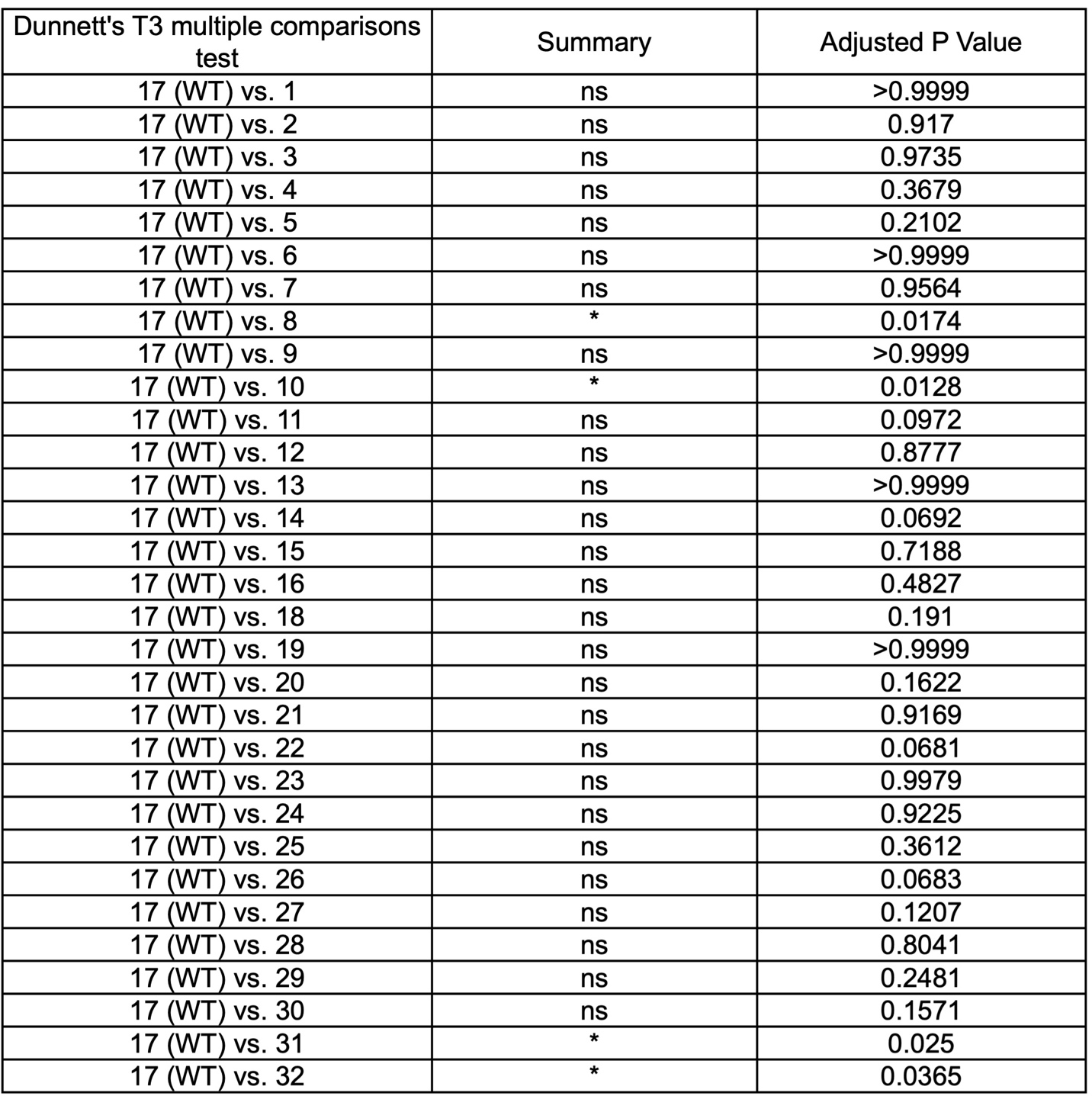
**Supplementary Table T6. Dunnett’s post hoc analysis for cellulose – I lysate screen data at pH 5.5.** Post hoc follow-up analysis after Welch’s ANOVA comparing the mean activity for each supercharged construct to the native enzyme. This statistical analysis corresponds to the data shown in **Figure 2B** of the main manuscript and **Figure S1B** of the SI appendix.

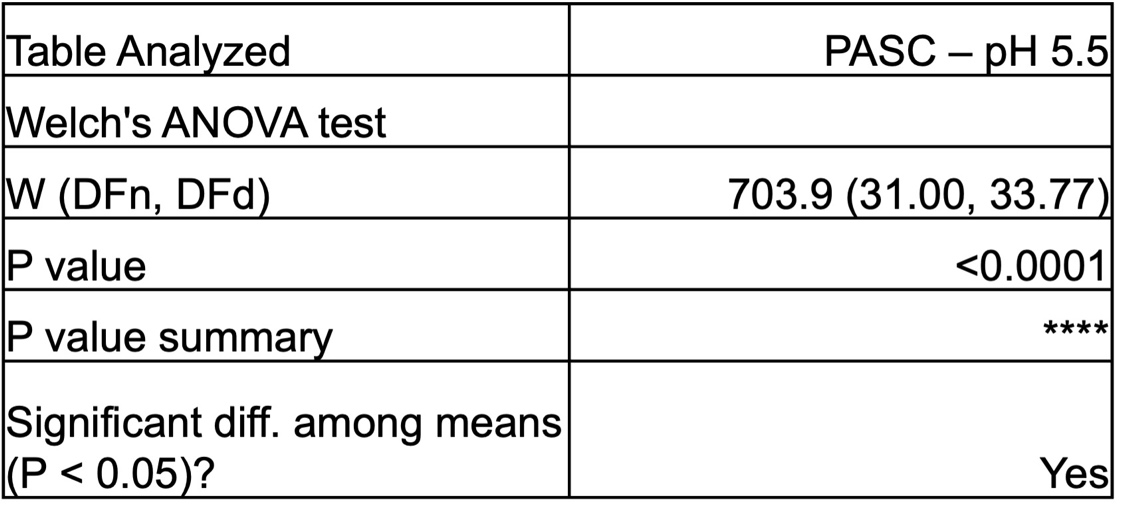

**Supplementary Table T7. Welch’s ANOVA result for PASC lysate screen data at pH 5.5.** Anova result depicted indicates a statistically significant difference (p ≤ 0.05) in mean activity for the supercharged library. This statistical analysis corresponds to the data shown in **Figure 2C** of the main manuscript.

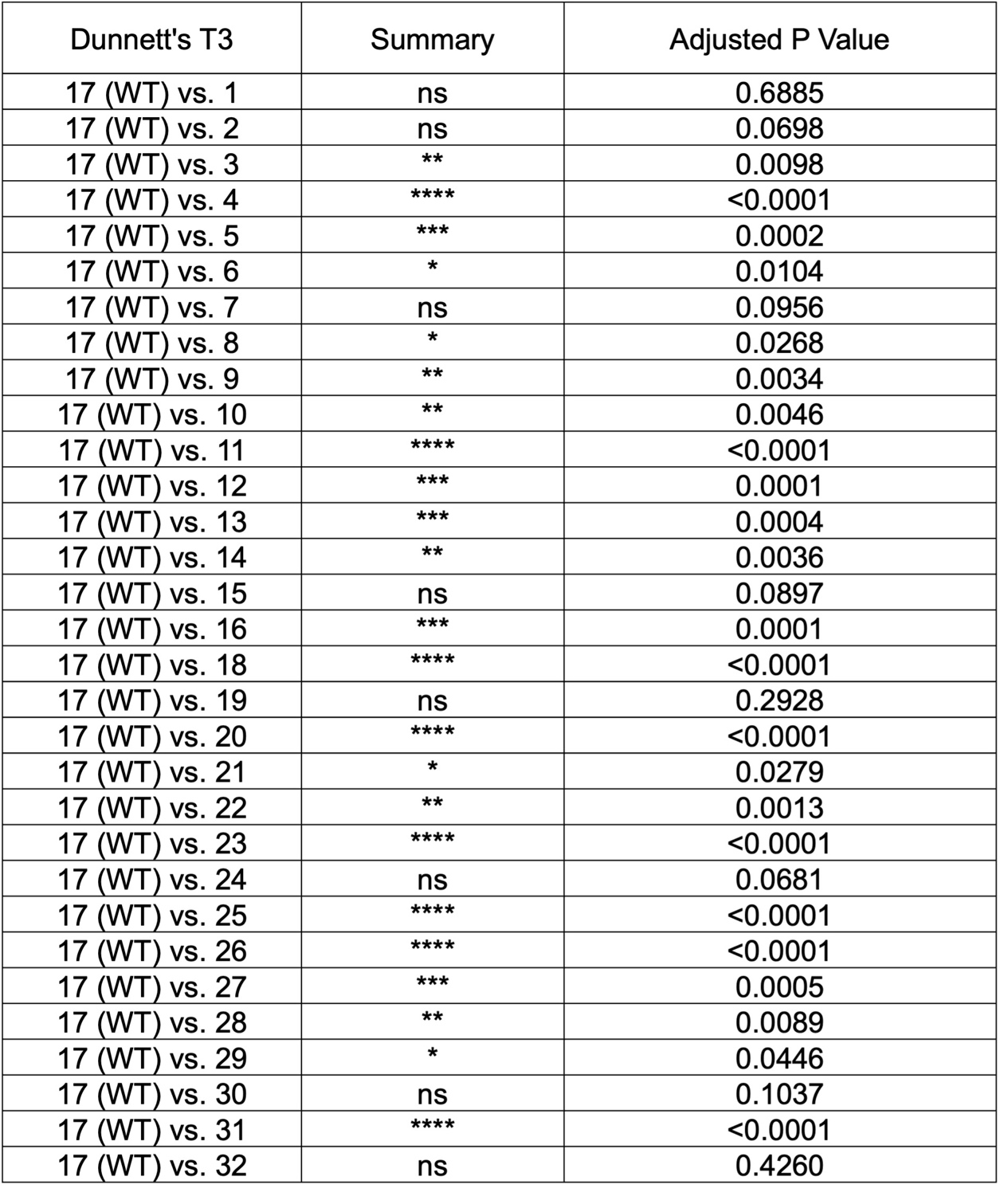

**Supplementary Table T8. Dunnett’s post hoc analysis for PASC lysate screen data at pH 5.5.** Post hoc follow-up analysis after Welch’s ANOVA comparing the mean activity for each supercharged construct to the native enzyme. This statistical analysis corresponds to the data shown in **Figure 2C** of the main manuscript.

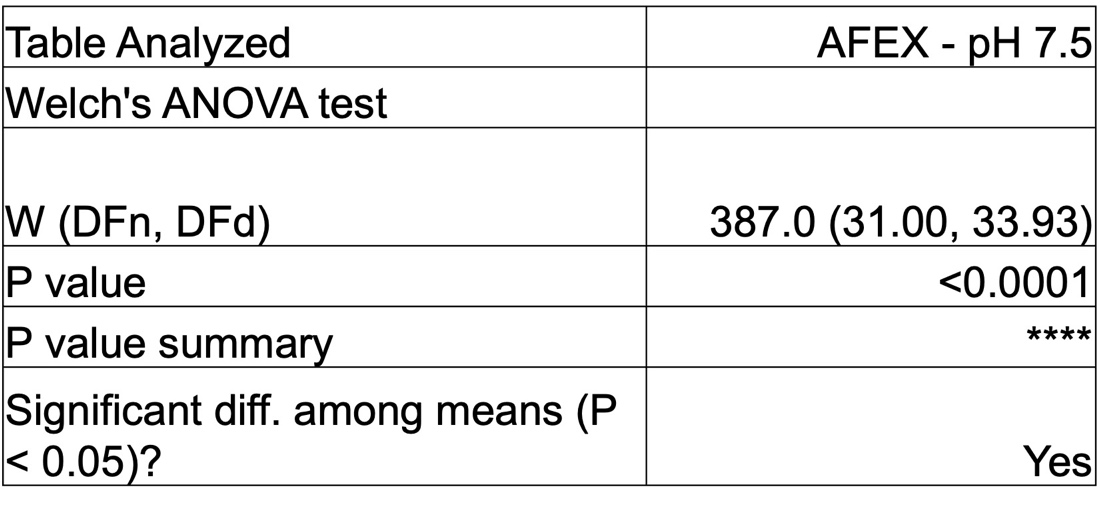

**Supplementary Table T9. Welch’s ANOVA result for AFEX corn stover lysate screen data at pH 7.5.** Anova result depicted indicates a statistically significant difference (p ≤ 0.05) in mean activity for the supercharged library. This statistical analysis corresponds to the data shown in **Figure 2D** of the main manuscript and **Figure S1C** of the SI appendix.

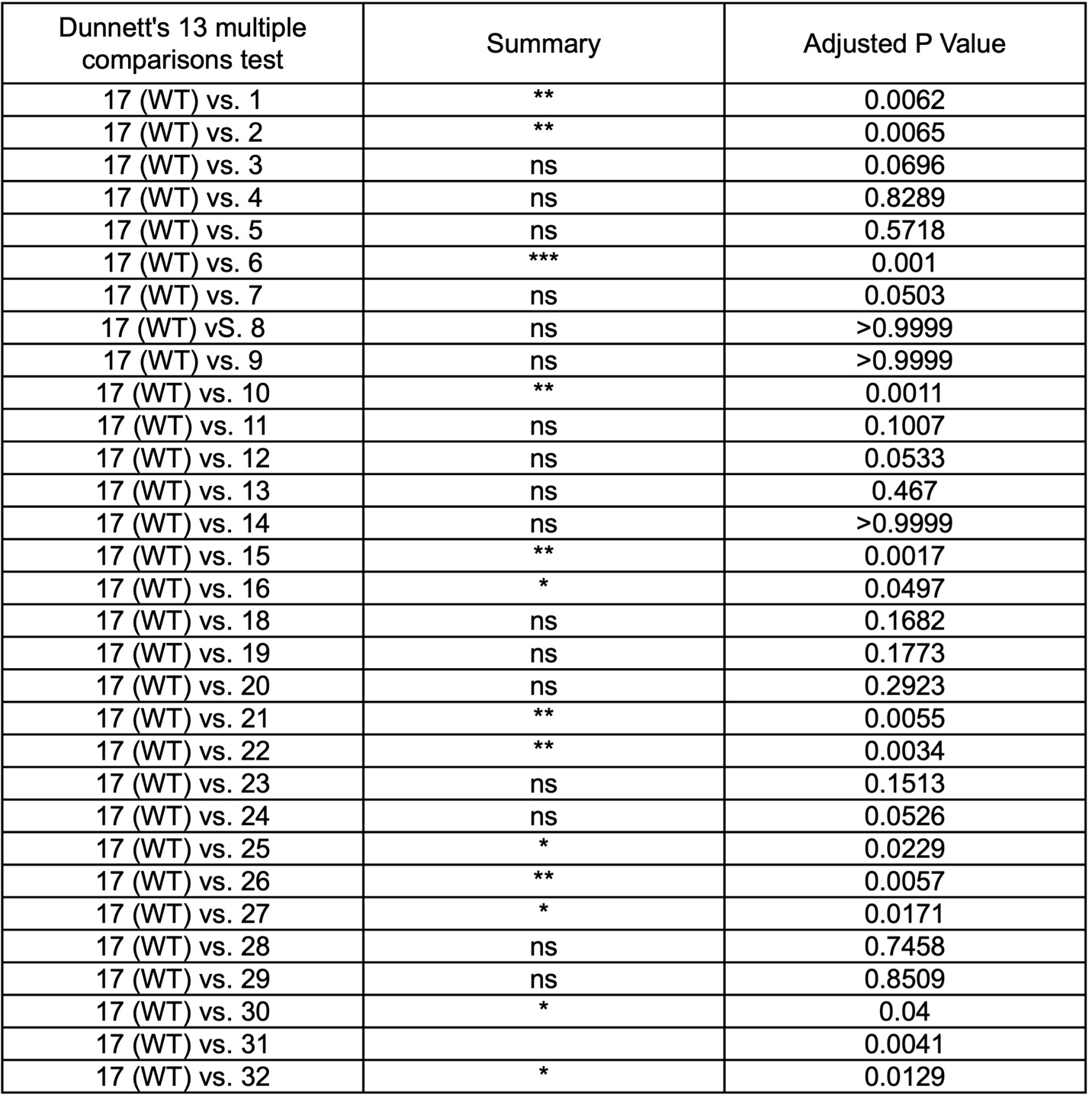
**Supplementary Table T10. Dunnett’s post hoc analysis for AFEX corn stover lysate screen data at pH 7.5.** Post hoc follow-up analysis after Welch’s ANOVA comparing the mean activity for each supercharged construct to the native enzyme. This statistical analysis corresponds to the data shown in **Figure 2D** of the main manuscript and **Figure S1C** of the SI appendix.

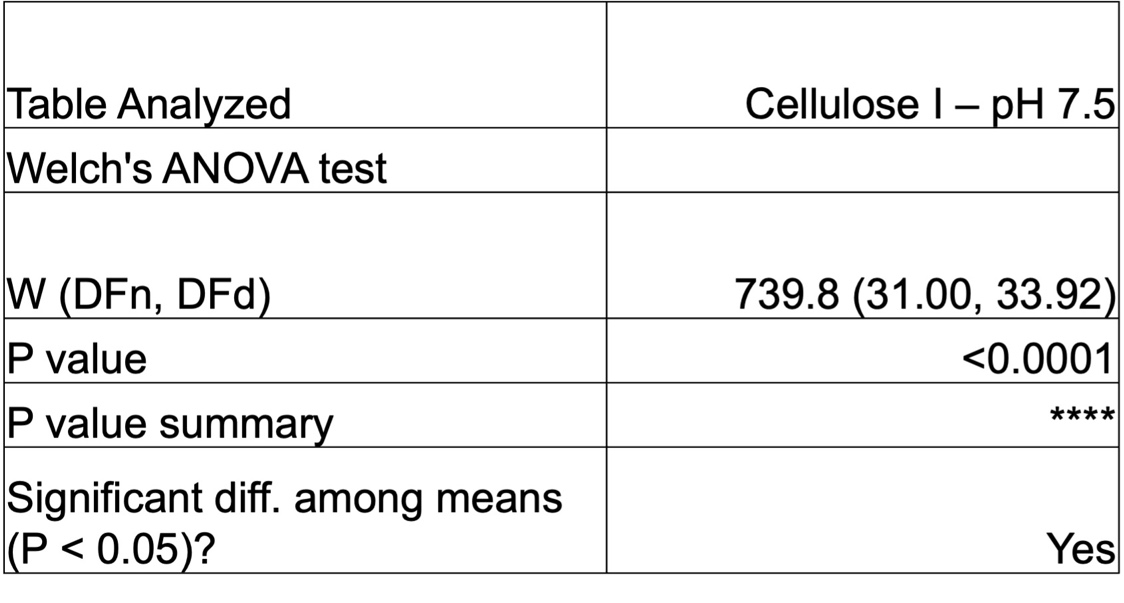

**Supplementary Table T11. Welch’s ANOVA result for cellulose – I lysate screen data at pH 7.5.** Anova result depicted indicates a statistically significant difference (p ≤ 0.05) in mean activity for the supercharged library. This statistical analysis corresponds to the data shown in **Figure 2E** of the main manuscript and **Figure S1D** of the SI appendix.

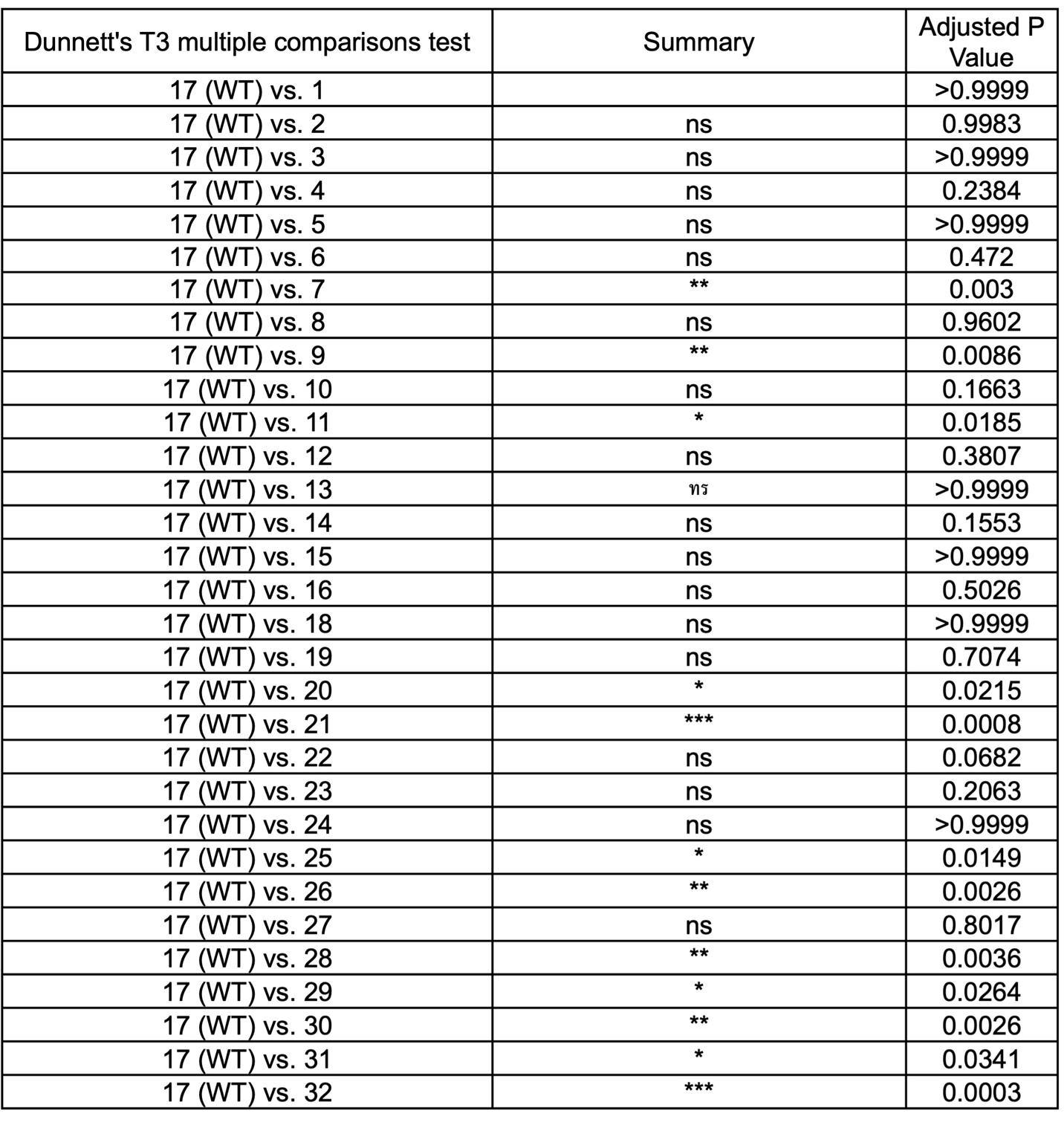
**Supplementary Table T12. Dunnett’s post hoc analysis for cellulose – I lysate screen data at pH 7.5.** Post hoc follow-up analysis after Welch’s ANOVA comparing the mean activity for each supercharged construct to the native enzyme. This statistical analysis corresponds to the data shown in **Figure 2E** of the main manuscript and **Figure S1D** of the SI appendix

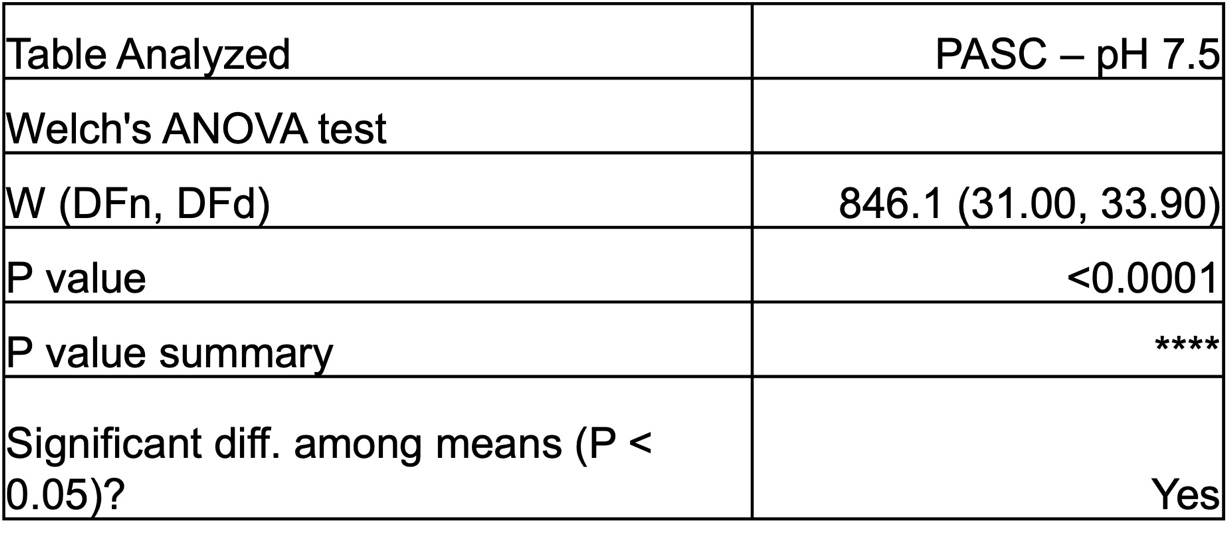

**Supplementary Table T13. Welch’s ANOVA result for PASC lysate screen data at pH 7.5.** Anova result depicted indicates a statistically significant difference (p ≤ 0.05) in mean activity for the supercharged library. This statistical analysis corresponds to the data shown in **Figure 2F** of the main manuscript.

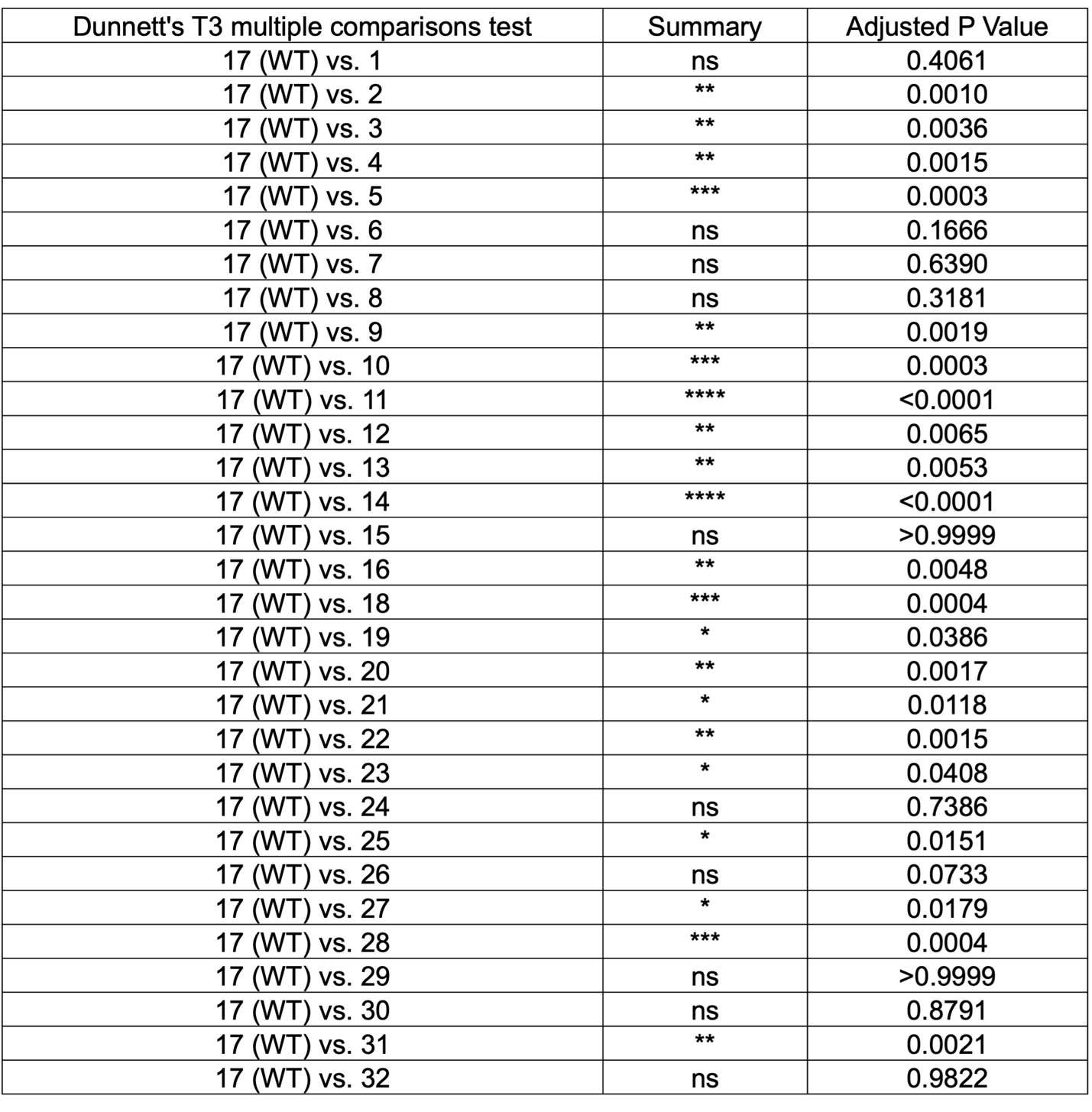
**Supplementary Table T14. Dunnett’s post hoc analysis for PASC lysate screen data at pH 7.5.** Post hoc follow-up analysis after Welch’s ANOVA comparing the mean activity for each supercharged construct to the native enzyme. This statistical analysis corresponds to the data shown in **Figure 2F** of the main manuscript.

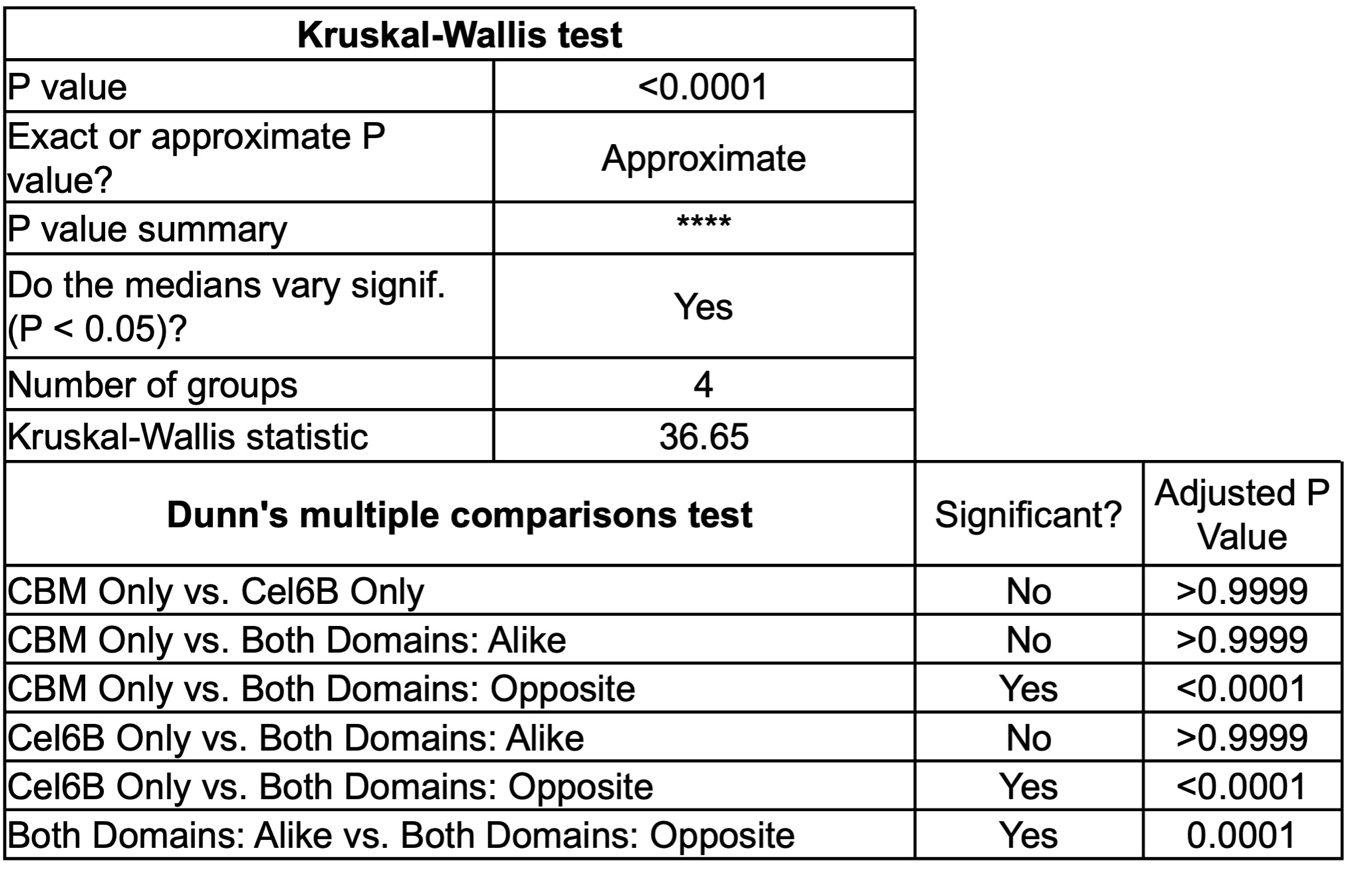
**Supplementary Table T15. Kruskal-Wallis test and Dunn’s post hoc analysis for comparing mutation placement to median activity on AFEX corn stover at pH 5.5.** ANOVA to determine if there is a significant difference in median activity for constructs containing only a supercharged CBM, Cel6B, two similar charged, or two oppositely charged domains. Dataset was non-parametric thus comparisons cannot be made on the mean and must analyze medians instead using Dunn’s post hoc analysis. This statistical analysis corresponds to the data shown in **Figure S2A** of the SI appendix.

**Supplementary Table T16. Kruskal-Wallis test and Dunn’s post hoc analysis for comparing mutation placement to median activity on AFEX corn stover at pH 7.5.** ANOVA to determine if there is a significant difference in median activity for constructs containing only a supercharged CBM, Cel6B, two similar charged, or two oppositely charged domains. Dataset was non-parametric thus comparisons cannot be made on the mean and must analyze medians instead using Dunn’s post hoc analysis. This statistical analysis corresponds to the data shown in **Figure S2B** of the SI appendix
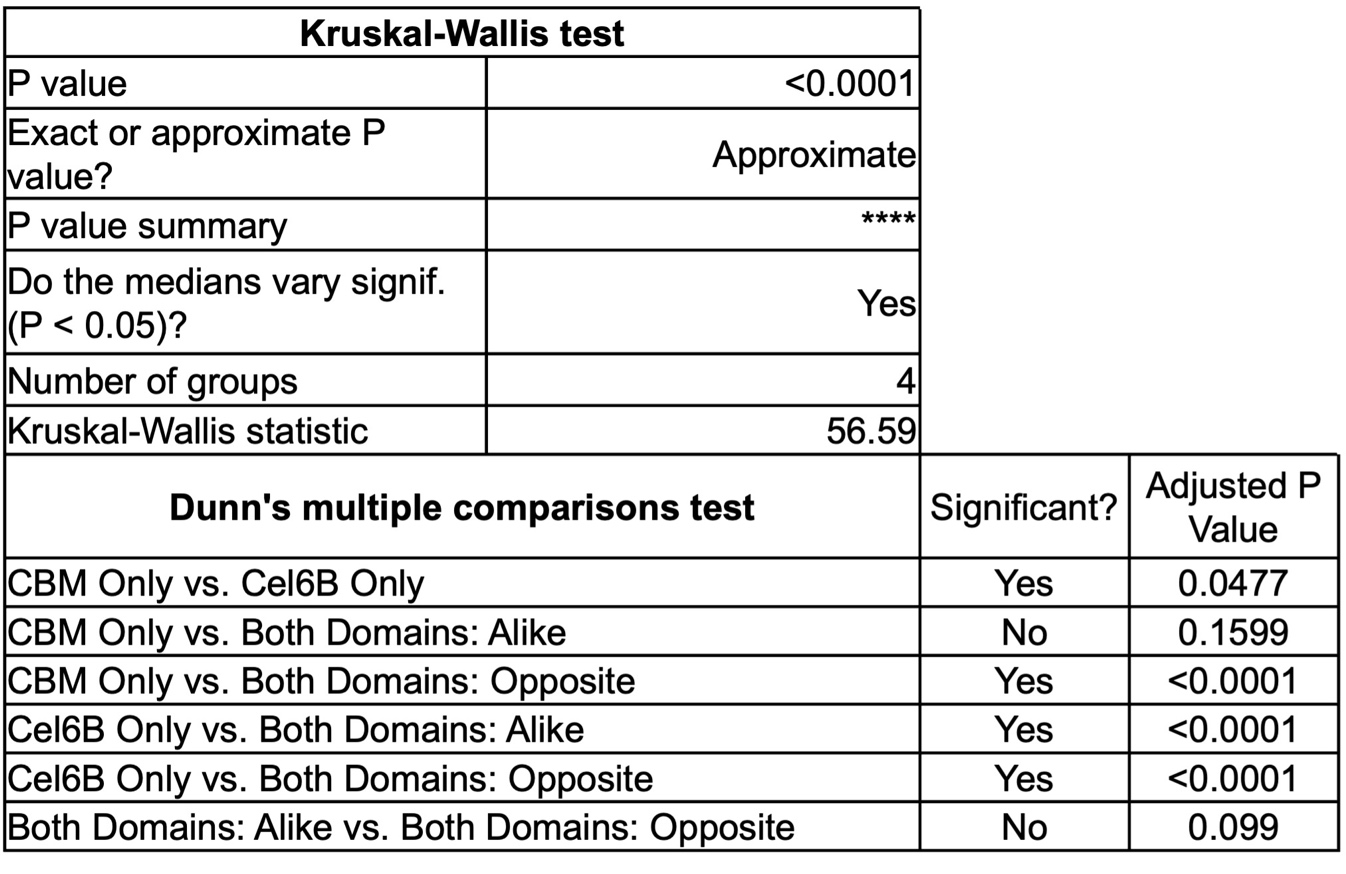
.

**Supplementary Table T17. Kruskal-Wallis test and Dunn’s post hoc analysis for comparing mutation placement to median activity on cellulose – I at pH 5.5.** ANOVA to determine if there is a significant difference in median activity for constructs containing only a supercharged CBM, Cel6B, two similar charged, or two oppositely charged domains. Dataset was non-parametric thus comparisons cannot be made on the mean and must analyze medians instead using Dunn’s post hoc analysis. This statistical analysis corresponds to the data shown in **Figure S3A** of the SI appendix
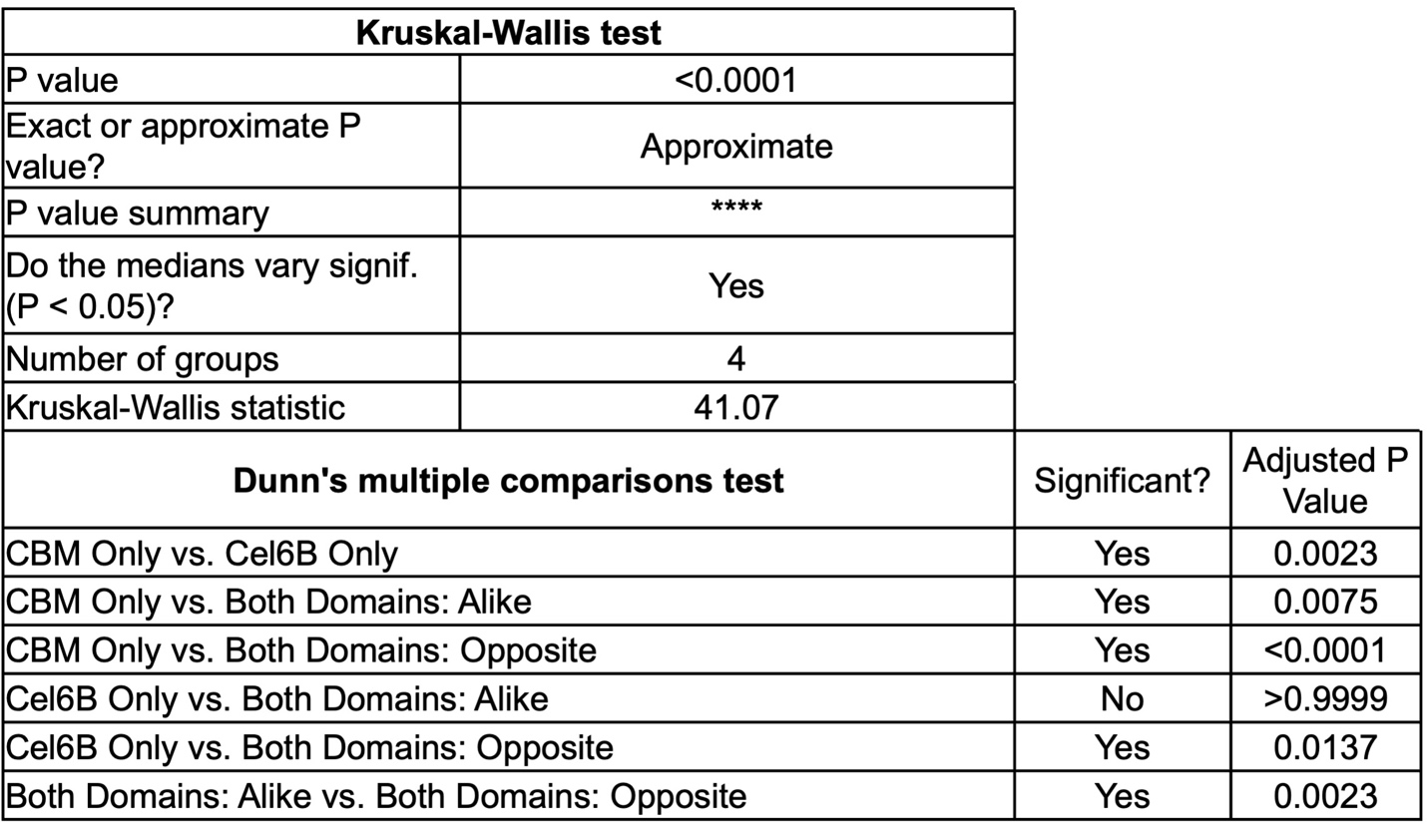
.

**Supplementary Table T18. Kruskal-Wallis test and Dunn’s post hoc analysis for comparing mutation placement to median activity on cellulose – I at pH 7.5.** ANOVA to determine if there is a significant difference in median activity for constructs containing only a supercharged CBM, Cel6B, two similar charged, or two oppositely charged domains. Dataset was non-parametric thus comparisons cannot be made on the mean and must analyze medians instead using Dunn’s post hoc analysis. This statistical analysis corresponds to the data shown in **Figure S3B** of the SI appendix
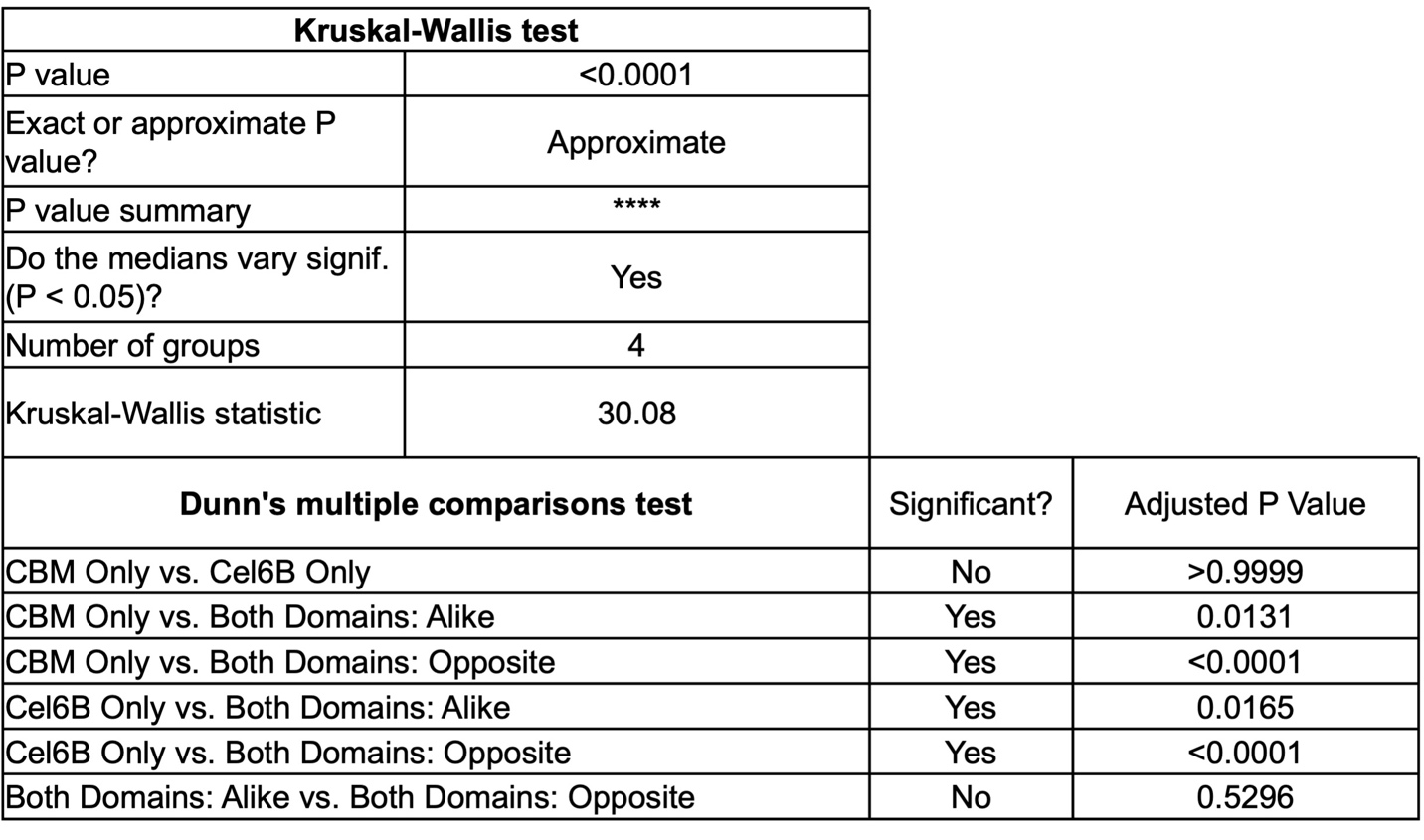
.
